# CORNICHON HOMOLOG 5-dependent ER export of membrane cargoes in phosphate-starved *Arabidopsis* root as revealed by membrane proteomic analysis

**DOI:** 10.1101/2024.12.30.630708

**Authors:** Tzu-Yin Liu, Ming-Hsuan Tsai, Jhih-Yi Wang, Hui-Fang Lung, Chiao-An Lu, Chang-Yi Chiu, Hong-Xuan Chow

## Abstract

Developing plants tolerant of low phosphate (Pi) availability is essential to reduce reliance on fertilizers and achieve agricultural sustainability. One strategy is to enhance the endoplasmic reticulum (ER) export of cargoes associated with Pi starvation and their trafficking to final destinations. However, the mechanisms underlying this process are underexplored. We recently discovered that *Arabidopsis thaliana CORNICHON HOMOLOG 5* (*AtCNIH5*) encodes a Pi deficiency-induced ER cargo receptor that regulates Pi homeostasis. To find potential membrane cargoes of *At*CNIH5, we applied the UV-cleavable 4-hexylphenylazosulfonate (Azo)-solubilized microsomal protein extraction for iTRAQ-based proteomic analysis. We identified 4,317 proteins in Pi-limited *Arabidopsis* roots, with 372 upregulated and 106 downregulated proteins in *cnih5*. Besides PHOSPHATE TRANSPORTER 1 proteins (PHT1s), downregulation of the biosynthetic or modifying enzymes for cell wall polysaccharides, very long-chain fatty acids, and their derivative extracellular aliphatic compounds is over-represented. Using the yeast split-ubiquitin and the *in-planta* tripartite split-GFP assays, we verified the interaction of *At*CNIH5 with various downregulated transporters in *cnih5*, including *At*PHT1s, *At*OCT1, *At*URGT6, *At*DTX21, and *At*DTX35. In addition, we demonstrated that the C-terminal acidic residue of *At*CNIH5 is required for interaction with *At*OCT1 but not with *At*PHT1;1 or *At*DTX21, indicating distinct cargo selection mechanisms. More importantly, enhancing *in-situ At*CNIH5 expression/activity enhances plant growth. By analogy with transcriptional factors that govern gene expression, we propose that *At*CNIH5 acts as a low Pi-responsive hub to facilitate ER export of specific membrane cargoes, providing a potential engineering strategy to improve plant fitness under suboptimal Pi supply.

## Introduction

Phosphorus (P) is an essential macronutrient for plant growth and development (Poirier and Bucher, 2002; Poirier et al., 2022). Due to the low concentration of inorganic phosphate (Pi) in soil, the primary form of P that plants absorb, applying Pi fertilizer has become a common practice to improve crop productivity. However, the limited mobility of Pi in soil prevents plants from acquiring most applied Pi fertilizer, leading to P runoff into aquatic ecosystems. Therefore, developing plants with increased Pi uptake and high-use efficiency is crucial for minimizing Pi fertilizer usage, thereby reducing environmental pollution and promoting agricultural sustainability.

To adapt to low Pi bioavailability, plants employ various strategies to enhance external Pi acquisition and internal Pi remobilization, involving distinct Pi transporters in distinct subcellular compartments. *PHOSPHATE TRANSPORTER 1* (*PHT1*) genes encode the plasma membrane (PM)-localized Pi transporters responsible for the uptake and the intercellular translocation of Pi. Even though Pi deprivation upregulates most *PHT1* genes at the transcript level, the translation, secretion, and recycling rate of PHT1 proteins (PHT1s) also determine their abundance at the PM (Nussaume et al., 2011). Multiple post-translational mechanisms have been elucidated to regulate the protein level and the PM localization of PHT1s. The plant-specific SEC12-related PHOSPHATE TRANSPORTER TRAFFIC FACILITATOR1 (PHF1) contributes to the accumulation of PHT1s at the PM by assisting the PHT1s transit through the endoplasmic reticulum (ER) (González *et al*., 2005; Bayle *et al*., 2011; Chen *et al*., 2015). The rice casein kinase II (CK2) diminishes the interaction between *Os*PT8 and *Os*PHF1 by phosphorylating *Os*PT8, thereby leading to ER retention of *Os*PTs (Chen et al., 2015). On the contrary, the protein phosphatase type 2C *Os*PP95 acts antagonistically with *Os*CK2 (Yang et al., 2020). In addition, under Pi sufficiency, the ubiquitin-conjugating E2 enzyme UBC24/PHOSPHATE2 (PHO2) at the ER or post-ER compartments mediates the ubiquitination of the endomembrane-associated *At*PHT1s followed by the vacuolar degradation (Huang et al., 2013). The PM-localized ubiquitin E3 ligase NITROGEN LIMITATION ADAPTATION (NLA) participates in the ubiquitination-mediated and clathrin-dependent endocytosis of PHT1s when Pi is replete (Lin *et al*., 2013; Park *et al*., 2014).

The ER is the start site where newly synthesized PHT1s travel towards the PM. The ER-to-Golgi transport of PHT1s relies on the coat protein complex II (COPII)-mediated vesicles or carriers (Bayle et al., 2011). However, it remains unknown how diverse membrane cargoes associated with Pi starvation are transported in the late secretory pathway and whether post-translational modification events regulate their packaging into COPII vesicles. The formation of COPII vesicles is a multi-step process that involves the recruitment of five core cytosolic factors, SAR1, SEC23, SEC24, SEC13, and SEC31, to the ER exit sites (ERES) (Barlowe and Miller, 2013; Kurokawa and Nakano, 2018). In particular, incorporation of multi-spanning transmembrane (TM) proteins into COPII vesicles in the budding yeast *Saccharomyces cerevisiae* requires dual binding with the COPII coat subunit Sec24 and the ER cargo receptor Erv14 (Powers and Barlowe, 1998; Harmel et al., 2012; Pagant et al., 2015). The role of Erv14 in COPII coupling is likely due to its function as a membrane chaperone, which exposes the sorting signals of Erv14 clients for their interaction with Sec24 and efficient capture into COPII vesicles (Dancourt and Barlowe, 2010; Pagant et al., 2015). Loss of the fungal Erv14 compromised the targeting of several transporters destined for the late secretory compartments, thus resulting in mis-regulating numerous cellular functions (Powers and Barlowe, 1998; Nakanishi et al., 2007; Pagant et al., 2015; Zimmermannová et al., 2019; Zheng et al., 2023). The *Drosophila* Cornichon (Cni), like its yeast homolog Erv14, was shown to recruit the epidermal growth factor (EGF)-like ligand Gurken into COPII vesicles (Roth *et al*., 1995; Bökel *et al*., 2006). Without Cni, the ER export and secretion of Gurken are disrupted (Bökel et al., 2006).

Similarly, the plant CORNICHON HOMOLOG proteins (CNIHs) act as the ER cargo receptors for various ion and metabolite transporters (Rosas-Santiago *et al*., 2015; Wudick *et al*., 2018; Wei *et al*., 2023; Yáñez-Domínguez *et al*., 2023). The *Arabidopsis thaliana* CNIHs (*At*CNIH1–5) have three putative transmembrane domains (TMDs), with the N terminus facing the cytosol, the C terminus facing the ER lumen, and at least four functional motifs (Wudick *et al*., 2018; Chiu *et al*., 2024). The cornichon motif is located between the first and second TMD of Erv14 and is well-conserved across all *At*CNIHs (Wudick *et al*., 2018). The FLN motif, known as a cargo-binding site in the second TMD of Erv14 (Pagant et al., 2015), and the cytosolic IFRTL motif, located between the second and third TMD of Erv14 and interacting with Sec24 (Powers and Barlowe, 2002), are also present in *At*CNIH2–4 with several residues changed. Lastly, the C-terminal acidic motif, only found in plants and fungi, is required for Erv14 to assist the PM targeting of the yeast Nha1, Pdr12, and Qdr2, as well as for the rice *Oryza sativa* CNIH1 (*Os*CNIH1) to aid the Golgi targeting of *Os*HKT1;3 (Rosas-Santiago et al., 2017).

We recently uncovered that *AtCNIH5* is a Pi starvation-induced (PSI) gene preferentially expressed in the outer layers of the root above the apical meristem (Chiu et al., 2024). *At*CNIH5 localizes to the ER and is closely associated with the *At*SAR1A/*At*SEC16A/*At*SEC24A-labeled ERES (Chiu et al., 2024). The T-DNA knockout mutant of *AtCNIH5* exhibited a lower abundance of *At*PHT1;1/2/3 and *At*PHT1;4 in the root microsomal fractions, with reduced shoot Pi levels and shoot-to-root distribution of ^32^P under Pi sufficiency and decreased ^32^P uptake under Pi deficiency (Chiu et al., 2024). The fluorescent protein tagged*-At*PHT1;1 in the Pi-limited root of *cnih5* was, in part, retained intracellularly in the root hair and in the epidermis within the transition/elongation zones (Chiu et al., 2024). Consistently, the fluorescence intensities of the *At*PHT1;1 fusion proteins diminished at the PM of the aforementioned root cells compared to the wild-type (WT) control (Chiu et al., 2024). These findings suggested a potential role for *At*CNIH5 in facilitating the transport of *At*PHT1s from the ER to the Golgi.

Intriguingly, phenotypes such as shorter root hairs and higher lateral root densities were also observed for *cnih5* (Chiu et al., 2024). Considering that *At*CNIH5 may serve as a cargo receptor for a broader range of membrane proteins beyond *At*PHT1s and that inefficient ER export of membrane cargoes in *cnih5* may cause their intracellular retention and eventual degradation due to the protein quality control, we reasoned that the steady-state levels of membrane cargoes of *At*CNIH5 would be reduced in *cnih5*. To identify potential cargoes of *At*CNIH5 associated with Pi starvation, we employed isobaric tags for relative and absolute quantification (iTRAQ)-based membrane proteomics. Because membrane proteins are highly hydrophobic and have low abundance, isolation of microsomal proteins (MPs) enriched for PM, ER, Golgi, vacuolar membranes, and endosomal vesicles is challenging (Wang and Liang, 2012). Conventionally, MPs are solubilized with sodium dodecyl sulfate (SDS) or other anionic detergents. However, SDS affects the performance of liquid chromatography-tandem mass spectrometry (LC–MS/MS), and removing SDS before mass spectrometry analysis results in an unavoidable loss of MPs (Wang and Liang, 2012). A study using the rice leaf as a model system demonstrated that the microcentrifugation-based microsomal protein extraction (MME) method is effective for downstream label-free quantitative proteome analysis, requiring neither a large amount of plant material nor ultracentrifugation to enrich PM proteins (Nguyen et al., 2021). The authors also applied the UV-cleavable and mass spectrometry-compatible surfactant 4-hexylphenylazosulfonate (Azo) (Brown *et al*., 2019; Brown *et al*., 2020) to minimize protein losses by omitting the SDS detergent wash step. Their results revealed several advantages of Azo over SDS in solubilizing the MPs, including a higher number and abundance of MPs (Nguyen et al., 2021). In this study, we modified the Azo-solubilized MME method to isolate root MPs from Pi-limited *Arabidopsis* WT and *cnih5* for iTRAQ proteomic analysis. We targeted our research to root MPs to reduce sample complexity, thus increasing the possibility of identifying functionally relevant protein changes in *cnih5*. Additionally, the membrane proteome analysis of Pi-limited roots provides valuable insights into the physiological role of *At*CNIH5. It also provides the first database of the membrane proteome for Pi-limited root biology research (Zhou et al., 2022).

With this approach, we identified 4,317 proteins in the Pi-limited WT and *cnih5* roots, in which 1,231 proteins (28.5%) have at least one TM helix. Of the differentially expressed proteins (DEPs) in *cnih5*, 372 and 106 proteins are upregulated and downregulated with ratios ≥ 1.25 or ≤ 0.75, respectively. Consistent with the primary role of *At*CNIH5 in the ER export of membrane cargoes, the Gene Ontology (GO) and Kyoto Encyclopedia of Genes and Genomes (KEGG) pathway enrichment analyses supported that the impairment of *At*CNIH5 leads to alterations in vesicular trafficking and suggested a close association of the downregulated proteins with protein export and localization. We also found that several transmembrane enzymes involved in the biosynthesis or modification of cell wall polysaccharides, very long-chain fatty acids (VLCFAs), and their derivative extracellular aliphatic compounds, such as cutin, suberin, and wax, are downregulated in Pi-limited *cnih5* roots. Using the yeast split-ubiquitin system (SUS) and the *in-planta* tripartite split-GFP assay, we confirmed the interaction of *At*CNIH5 with various potential membrane cargoes, including a number of transporters such as *At*PHT1s, *At*OCT1, *At*URGT6, *At*DTX21, and *At*DTX35. We generated a series of C-terminally truncated forms of *At*CNIH5 to find out which motif of *At*CNIH5 interacts with *At*PHT1;1. *At*CNIH5 1–26, which contains only the first TMD, is sufficient for localization to the ER in agro-infiltrated *Nicotiana benthamiana* (*N. benthamiana*) leaves and for interacting with *At*PHT1;1 in plant and yeast cells. Interestingly, the C-terminal tail of *At*CNIH5, which carries one aspartic acid residue, is required for interaction with *At*OCT1 but not with *At*PHT1;1 or *At*DTX21. Therefore, unlike the fungal and rice CNIHs interacting with their cognate cargoes through the C-terminal acidic motif, *At*CNIH5 employs a distinct mechanism to select *At*PHT1;1 as its cargo. Importantly, introducing the *At*CNIH5 genomic-GFP translation fusion to complement the *cnih5* mutant boosted plant growth under suboptimal Pi supply, suggesting that *At*CNIH5 acts as a hub to facilitate the ER export of specific membrane cargoes associated with Pi starvation to improve plant fitness.

## Results

### Azo-solubilized microcentrifugation-based microsomal protein extraction (MME) of *Arabidopsis* roots

Because the application of Azo promoted fast and reproducible tryptic digestion for quantitative proteomics (Brown et al., 2020), we replaced the mass spectrometry-incompatible surfactant SDS with Azo and established an efficient workflow of MPs extraction for quantitative proteomic analysis of *Arabidopsis* roots (**Fig. 1A**). To ensure that the Azo wash step can be skipped prior to mass spectrometry analysis, we tested the time-course effect of UV irradiation on Azo breakdown (Brown et al., 2019). Within 40 min of UV illumination, 1% Azo in 100 mM triethylammonium bicarbonate (TEABC) almost wholly degraded (**Fig. 1B**). After tryptic digestion and Azo degradation, small aliquots of four samples—two biological replicates each for WT and *cnih5*—were pre-analyzed in a single one-dimensional (1D)–LC–MS/MS run. A summary of the label-free proteomic analysis for quality check is shown in Fig. S1A. We used multiple marker abundance profiling (MMAP) (Hooper et al., 2016) to estimate the relative abundance of organelles in the MPs samples. The results suggested that the modified Azo-solubilized MME method greatly removed proteins targeted to the plastid, nucleus, and extracellular space while enriching proteins targeted to the mitochondria, ER, Golgi, PM, and vacuole (Fig. S1B, C). To further evaluate this modified MME method, we examined the abundance of the well-established root MPs, including the PM-targeted *At*NRT2.1 (Chopin et al., 2007), *At*PHT1;1/2/3 and *At*PHT1;4, the Golgi-localized *At*PHO1 (Arpat *et al*., 2012; Liu *et al*., 2012), and the ER-resident *At*PHF1. While the amount of NRT2.1 declined under Pi starvation and showed no differences between WT and *cnih5*, the levels of *At*PHT1;1/2/3, *At*PHT1;4, *At*PHF1, and *At*PHO1 were upregulated following Pi deprivation (**Fig. 2A**), as previously reported. Importantly, the abundance of *At*PHT1;1/2/3 and *At*PHT1;4 decreased in *cnih5* compared to the WT controls, regardless of Pi availability, whereas *At*PHF1 proteins were more abundant in *cnih5* than in WT (**Fig. 2A**), which aligns with our recent immunoblot results obtained from commercial membrane protein extraction or total protein extraction methods (Chiu et al., 2024). Collectively, these data justified the suitability of the modified Azo-solubilized MME method for enriching root MPs.

**Fig 1.**
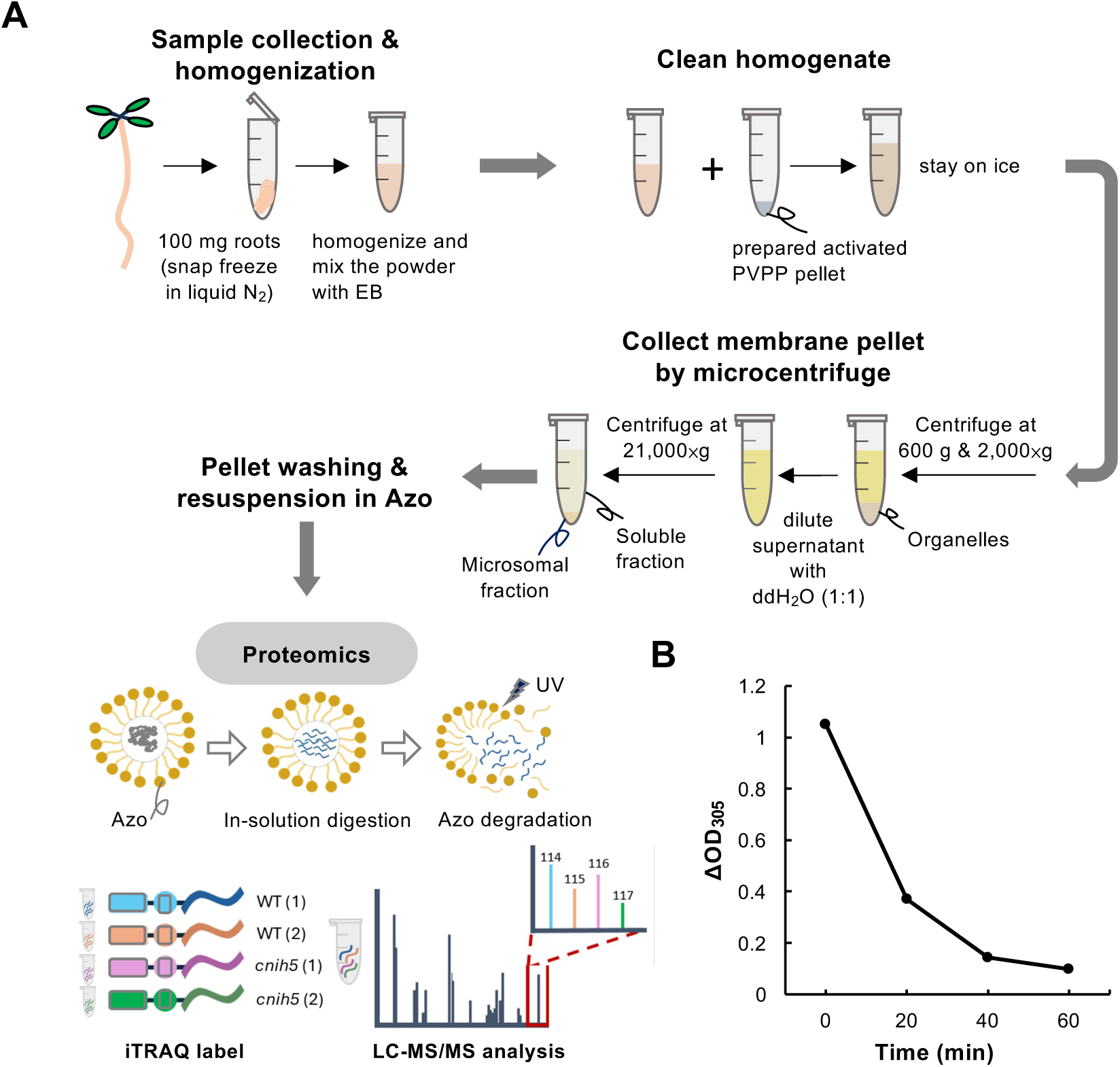
iTRAQ-based quantitative proteomic analysis of Pi-limited *Arabidopsis* roots. (A) Schematic workflow of Azo-solubilized microsomal protein extraction for iTRAQ-based quantitative proteomic analysis of *Arabidopsis* Pi-limited roots. (B) UV-Vis spectrum monitoring the degradation of 1% Azo as a function of UV irradiation time.

**Fig 2.**
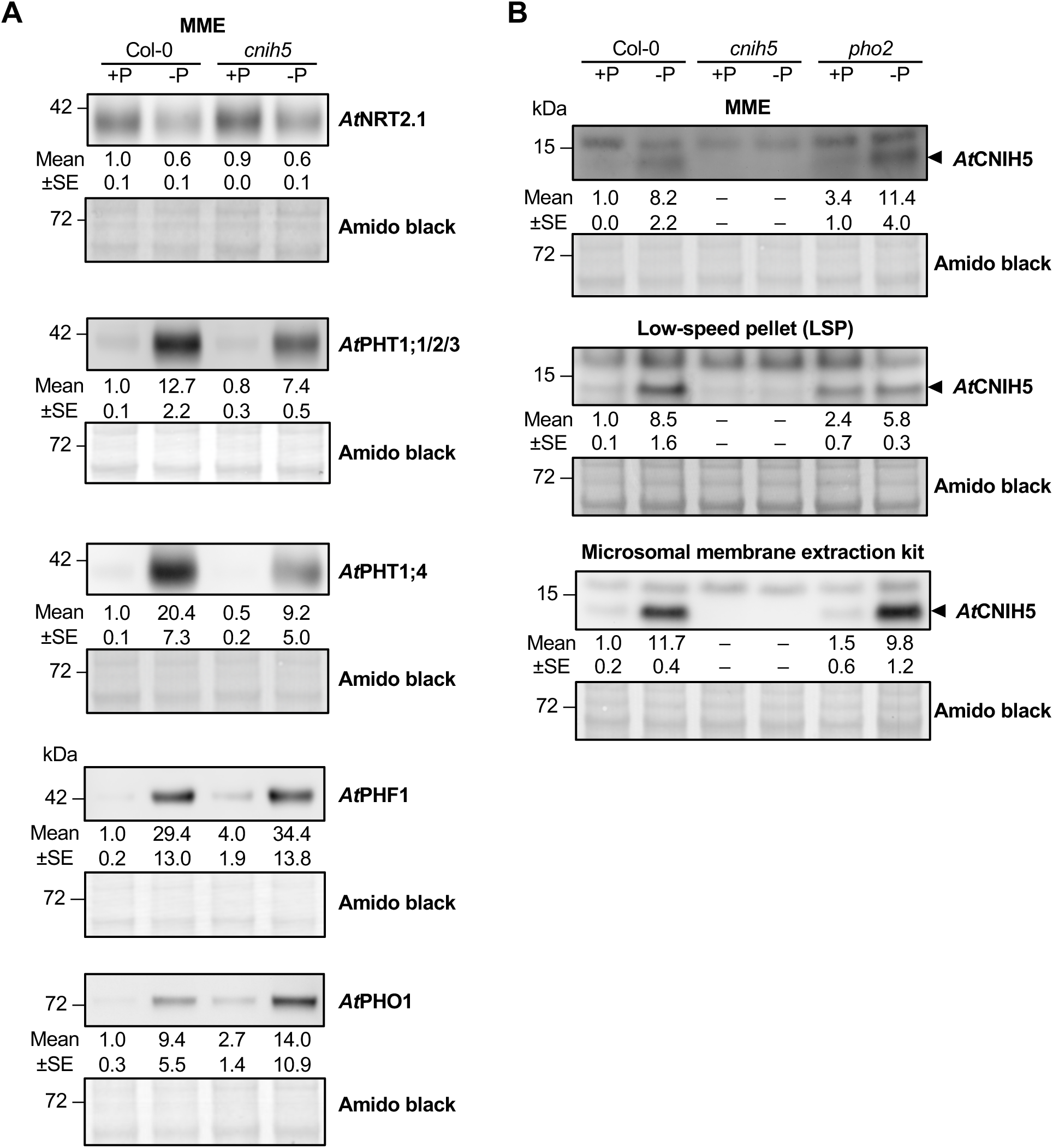
Immunoblot analysis of root microsomal proteins extracted from WT, *cnih5*, and *pho2* mutants. (A, B) Protein expression of *At*NRT2.1, *At*PHT1;1/2/3, *At*PHT1;4, *At*PHF1, and *At*PHO1 (A) and *At*CNIH5 (B) in the root of 11-day-old seedlings of *Arabidopsis* WT, *cnih5*, and *pho2* under +P (250 µM KH_2_PO_4_) and –P (0 µM KH_2_PO_4_, five days of starvation) conditions. The band of *At*CNIH5 is indicated by arrowhead. The relative protein expression level was normalized with the corresponding amido black staining and relative to the WT control. Error bars represent SE (n = 2–3). The applied extraction methods were shown as indicated. MME, microsomal protein extraction.

### Upregulation of *At*CNIH5 in the root during Pi starvation and in the Pi overaccumulator *pho2* under Pi sufficiency

Since *At*CNIH5 was not identified in our small-scale label-free proteomic analysis, we raised a specific polyclonal antibody to detect the abundance of *At*CNIH5 in the root MPs. This antibody recognized a band in the Pi-limited roots of WT seedlings at ∼15 kDa that was absent in *cnih5* (**Fig. 2B**). In agreement with the finding that low Pi induces the gene expression of *AtCNIH5* (Chiu et al., 2024), the protein amount of *At*CNIH5 was also elevated during Pi starvation (**Fig. 2B**), suggesting that *AtCNIH5* is upregulated by Pi starvation at the transcript and protein level. Of note, under Pi sufficiency, *At*CNIH5 modestly accumulated in the root of *pho2* relative to the WT control (**Fig. 2B**). This increase was recapitulated by two other different MPs extraction methods (**Fig. 2B**), validating the previous membrane proteomic analysis showing that the protein level of *At*CNIH5 significantly increased in the Pi-replete root of *pho2* relative to the WT (Huang et al., 2013). These results suggest that *At*CNIH5 expression is induced to meet the increased demand for ER export of *At*PHT1s, supporting the genetic evidence from our side-by-side study, which shows that loss of *AtCNIH5* alleviates the Pi toxicity of *pho2* (Chiu et al., 2024).

### Dysfunction of *At*CNIH5 alters vesicular trafficking, acyl-lipid metabolism, and cell wall composition under Pi limitation

After labeling the tryptic-digested peptides with iTRAQ reagents, we mixed and separated them using two-dimensional high-performance liquid chromatography (2D–HPLC). Each fraction was diluted and trapped onto a reverse-phase column for separation, followed by Orbitrap mass spectrometry. By combining the total proteins of the four samples, we identified 32,812 peptides and 21,433 unique peptides that matched with 4,317 proteins with confidence (criteria described in Materials and Methods), resulting in an average sequence coverage of 17.8% (Table S1). About 99.3% of the identified proteins were shared across the four samples (Table S1), indicating that our proteomic data were unbiased, as nearly all proteins within the detection range could be identified. The proteins containing TM helices were predicted or annotated by DeepTMHMM and UniProtKB (**Fig. 3A)** (Krogh *et al*., 2001; Hallgren *et al*., 2022; The UniProt Consortium, 2023). The 1,231 identified proteins (28.5%) have at least one TM helix and thus are integral membrane proteins (**Fig. 3A**). The subcellular localizations of the 4,317 identified proteins based on SUBA5 (Hooper et al., 2016) showed that 7.8% localize to the ER, 10.1% to the Golgi, 4.4% to the vacuole, and 16.2% to the PM (**Fig. 3B**). Consequently, about 37.3% (1,609) and 21.2% (915) represent MPs and integral MPs, respectively (**Fig. 3A**), while 19.3% of the 915 integral MPs localized to the ER, 15.7% to the Golgi, 9.7% to the vacuole, and 31.7% to the PM (**Fig. 3C**). The general and detailed descriptions of proteomics data, including the summary of proteomic analysis, protein scores and mass, annotation, and quantification for the identified proteins, are listed in Table S1.

**Fig 3.**
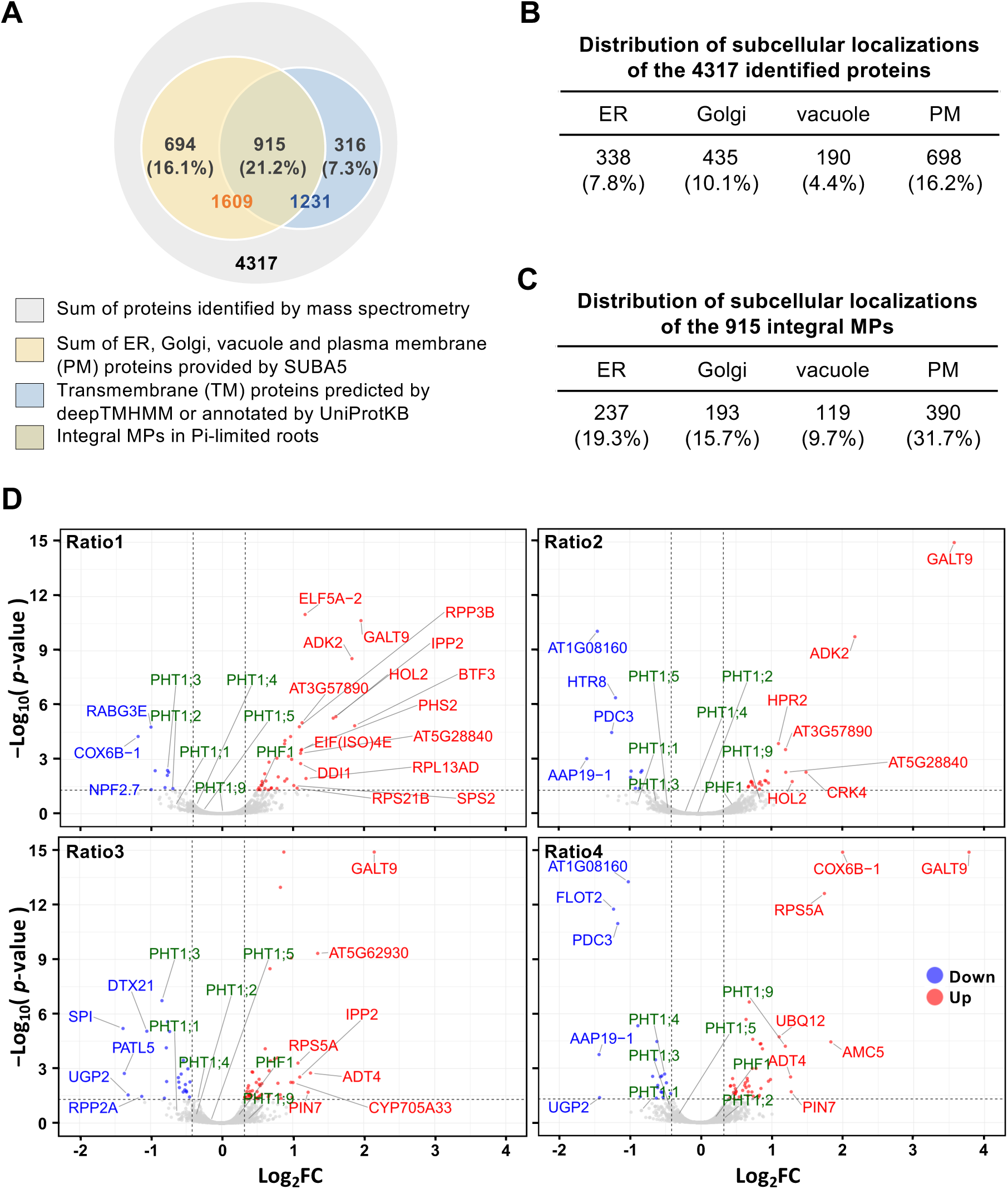
Summary of iTRAQ-based proteomic analysis of microsomal proteins (MPs) in Pi-limited WT and *cnih5* roots. (A) Venn diagram for comparing MPs and transmembrane (TM) proteins of the 4,317 identified proteins. Subcellular localization was analyzed by SUBA5. TM protein prediction or annotation was performed by deepTMHMM or UniprotKB. (B) Distribution of subcellular localizations of the identified 4,317 proteins analyzed by SUBA5. Note that not all proteins are restricted to a single subcellular location. (C) Distribution of subcellular localizations of the 915 integral MPs among the identified 1,231 TM proteins analyzed by SUBA5. Note that not all proteins are restricted to a single subcellular location. (D) Volcano plots of differentially expressed proteins (DEPs) in the Pi-limited root of *cnih5*. Results of the four comparison groups (four ratios of *cnih5* versus WT) were shown separately, with the −log_10_FDR adj. *p*-value plotted against the log_2_ fold-change (FC). The vertical dash lines denote ±1.25-FC (Log_2_ FC ≈ 0.322 and Log_2_FC ≈ −0.415), while the horizontal dash line denotes FDR adj. *p*-value = 0.05. Upregulated and downregulated proteins are, respectively, indicated by red and blue dots. Only DEPs with ±2-FC are marked in red or blue. The *At*PHT1s and *At*PHF1 are marked in green.

The 4,317 identified proteins were subjected to GO enrichment analysis by ShinyGO 0.82 (Ge et al., 2019). Only the top significant GO terms (FDR adj. *p*-value < 0.05) were shown according to the ranking of fold enrichment. The top terms are nucleoside-triphosphatase (NTPase) activity, pyrophosphatase (PPase) activity, and mRNA binding in the molecular function (MF) category; intracellular transport, protein transport, and establishment of protein localization in the biological process (BP) category; plant-type vacuole and cell-cell junction in the cellular component (CC) category (Table S1 and Fig. S2). These results indicated that Pi depletion significantly impacts NTP- and PPi-dependent cellular activities required for endomembrane trafficking.

In comparison to the previous finding that analyzed quantitative total protein profiling in Pi-deficient versus Pi-sufficient *Arabidopsis* roots (Lan et al., 2012), we observed that among the 209 significantly up-regulated proteins in their datasets (Pi-deficient/Pi-sufficient plants ratio of ≥ 1.29), 57 proteins—13 of which are integral MPs—overlapped with our identified 4,317 proteins in Pi-limited roots (Table S1). This overlap accounts for 27.3% of the PSI proteins in their study. By contrast, we identified an additional 902 integral MPs (Table S1), indicating that enriching MPs for proteomics greatly expands the inventory of identified proteins implicated in Pi starvation responses.

Consistent with the immunoblot results (**Fig. 2A**), our proteomic data showed decreased protein abundance of several *At*PHT1 members and increased protein abundance of *At*PHF1 in Pi-limited root of *cnih5* compared to WT (**Table 1**). Specifically, we detected downregulation of *At*PHT1;2, *At*PHT1;3, and *At*PHT1;4 in at least three ratios (fold changes), with at least one ratio having FDR adj. *p*-value ≤ 0.05. Although not statistically significant, the downregulation of *At*PHT1;1 and *At*PHT1;5, along with the upregulation of *At*PHF1, displayed a consistent trend across four ratios. In contrast, *At*PHT1;9 was upregulated in the Pi-limited root of *cnih5* (**Table 1**). Volcano plots summarized the results from the four comparisons and highlighted the differentially expressed *At*PHT1 members and *At*PHF1 (**Fig. 3D**).

**Table 1.** Summary of *At*PHT1s and *At*PHF1 abundance in Pi-limited WT and *cnih5* roots. The normalized ratios compared the protein expression in Col-0_1, Col-0_2, *cnih5*_1 and *cnih5*_2 samples. The FDR adj. *p*-values are in parentheses.

| Accession | Gene name | AGI code | Coverage<br>[%] | Unique<br>peptides | *cnih5*_1<br>/Col-0_1 | *cnih5*_2<br>/Col-0_1 | *cnih5*_1<br>/Col-0_2 | *cnih5*_2<br>/Col-0_2 |
| --- | --- | --- | --- | --- | --- | --- | --- | --- |
| Q8VYM2 | *AtPHT1;1* | AT5G43350 | 38 | 2 | 0.63<br>(0.41) | 0.54<br>(0.28) | 0.65<br>(0.31) | 0.56<br>(0.22) |
| Q96243 | *AtPHT1;2* | AT5G43370 | 30 | 1 | 0.61<br>(0.05) | 0.97<br>(1.00) | 0.78<br>(0.47) | 1.24<br>(0.67) |
| O48639 | *AtPHT1;3* | AT5G43360 | 10 | 2 | 0.58<br>(0.00) | 0.65<br>(0.24) | 0.56<br>(0.00) | 0.74<br>(0.14) |
| Q96303 | *AtPHT1;4* | AT2G38940 | 12 | 6 | 0.77<br>(0.40) | 0.86<br>(0.89) | 0.79<br>(0.06) | 0.70<br>(0.00) |
| Q8GYF4 | *AtPHT1;5* | AT2G32830 | 6 | 2 | 0.82<br>(0.78) | 0.73<br>(0.58) | 0.90<br>(0.87) | 0.80<br>(0.42) |
| Q9S735 | *AtPHT1;9* | AT1G76430 | 5 | 2 | 1.00<br>(0.99) | 1.79<br>(0.20) | 1.27<br>(0.79) | 2.88<br>(0.00) |
| Q8GYE0 | *AtPHF1* | AT3G52190 | 40 | 13 | 1.45<br>(0.05) | 1.37<br>(0.36) | 1.16<br>(0.48) | 1.14<br>(0.62) |

Coverage [%]: The percentage of the protein sequences covered by identified peptides.
Unique peptides: The total number of distinct peptide sequences unique to the protein group.

Although *At*PHT1;1 is a *bona fide* membrane cargo of *cnih5* (Chiu et al., 2024), we noticed that the ratios of quantified *At*PHT1s varied across the four comparisons (**Table 1**), likely due to the limited number of biological replicates. To generate a more comprehensive list of DEPs for GO enrichment analysis, we adjusted the stringency of the protein profile comparisons by defining DEPs as those with at least two ratios ≥ 1.25 or ≤ 0.75 without applying FDR-adjusted *p*-value cutoffs. We also excluded candidates showing any opposite trends (≥ 1.25 or ≤ 0.75) across the four comparisons. As a result, we identified 372 upregulated proteins and 106 downregulated proteins in *cnih5* (Table S2). The GO enrichment analysis (FDR adj. *p*-value < 0.05) for the 372 upregulated proteins revealed that the top terms include copper ion binding, mRNA binding, and oxidoreductase activity in the MF category; sulfur compound biosynthetic process and purine nucleotide metabolic process in the BP category; cell-cell junction and plasmodesma in the CC category (Table S2 and Fig. S3A). The KEGG pathway enrichment analysis indicated that the most significantly enriched pathways are ascorbate and aldarate metabolism, as well as fructose and mannose metabolism (Fig. S3B). These results indicated that the upregulated proteins in the Pi-limited *cnih5* roots encode several enzymes involved in carbohydrate metabolic pathways, which may help protect plant cells from oxidative damage, likely reflecting compensatory mechanisms for the loss of *At*CNIH5. Interestingly, when focusing on the 51 upregulated integral MPs, we found significant GO terms related to transporter activities (Fig. S3C) and KEGG analyses revealed a strong association with SNARE-mediated vesicular transport (Fig. S3D).

For GO enrichment analysis of the 106 downregulated proteins, the most significant GO terms within the MF category include 7S RNA binding and fatty acid elongase activity (**Fig. 4A** and Fig. S4A); within the BP categories, the top terms are protein transport, cellular macromolecule localization, and establishment of protein localization (**Fig. 4B** and Fig. S4A); within the CC categories, the enriched terms are the AP-type membrane coat adaptor complex (**Fig. 4C** and Fig. S4A). These results can be attributed to the downregulation of membrane protein localization, Golgi-associated clathrin assembly, and the signal recognition particle (SRP) subunits that mediate co-translational membrane protein targeting (Table S2). We thus surmised that the loss of *At*CNIH5 triggers negative feedback regulation of protein translation and the formation of clathrin-coated vesicles, thereby reducing futile coated-vesicle assembly and preventing the toxic accumulation of *At*CNIH5-dependent cargoes in the ER. Furthermore, the KEGG enrichment analysis indicated that the downregulated proteins were associated with processes such as fatty acid elongation, protein export, cutin/suberin/wax biosynthesis, and modification of cell wall polysaccharides (Fig. S4B). Overall, the downregulated proteins in Pi-limited *cnih5* roots are linked to changes in vesicular trafficking, acyl lipid metabolism, and cell wall composition. Because of the limited availability of other antibodies for further validation of our quantitative proteomic data by immunoblotting, we conducted an additional label-free membrane proteomics on Pi-limited roots of WT and *cnih5* across three biological replicates. Label-free proteomics without fractionation yielded a dataset of 2035 identified proteins, which is smaller than our iTRAQ-based dataset (Table S1). Nonetheless, we found an overlap of 29 proteins between the label-free dataset and the iTRAQ-label downregulated proteins in Pi-limited *cnih5* roots (Table S1) and validated that *At*PHT1;1, *At*PHT1;2, *At*PHT1;3, *At*PHT1;4, *At*EXPA17, *At*GXM2, *At*AAP19-1, and *At*GAE1 showed a similar trend of downregulation (normalized ratios ≤ 0.82; Table S3), supporting that these proteins are truly downregulated in Pi-limited *cnih5* roots.

**Fig 4.**
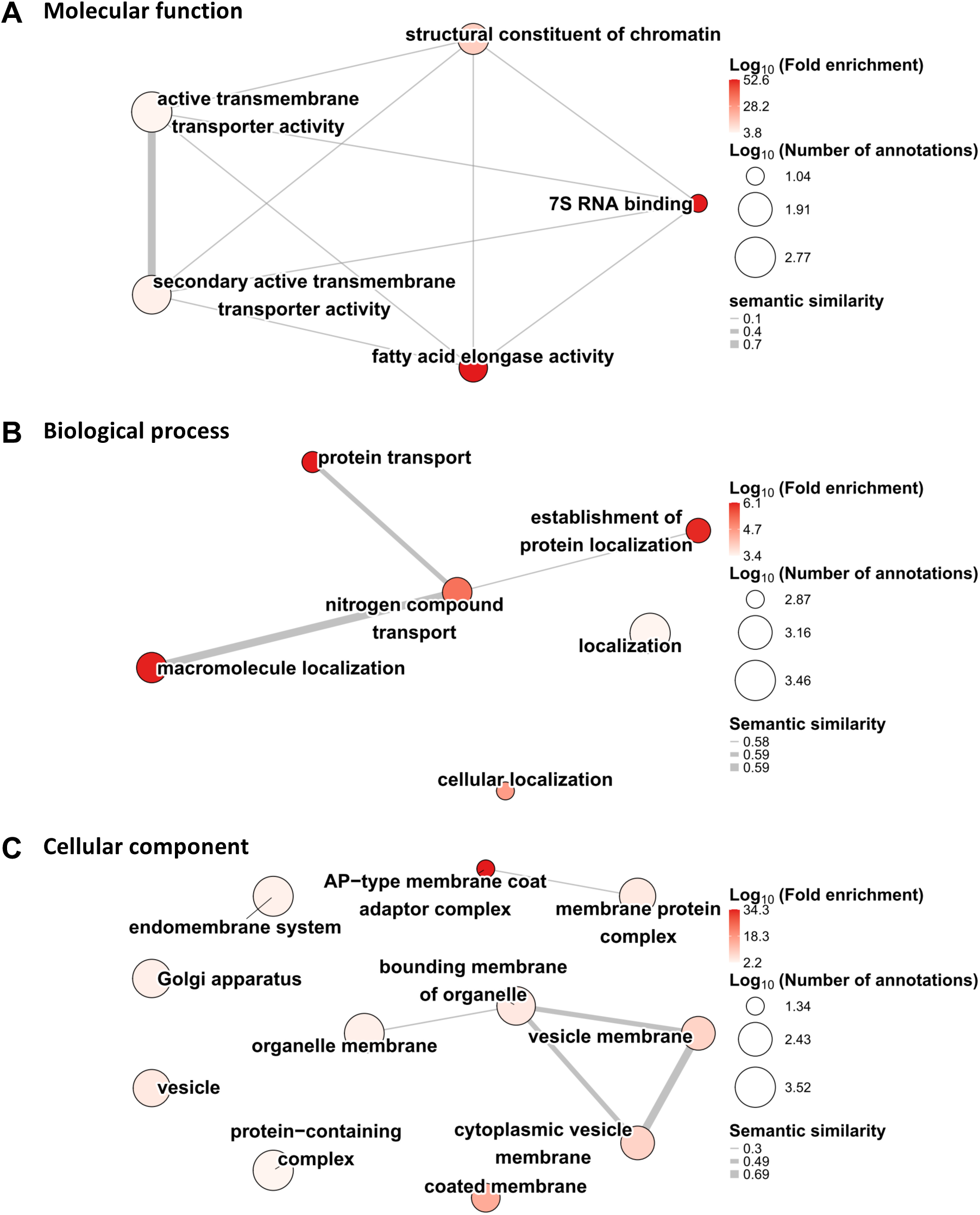
Interactive graph of enriched GO terms and semantic similarities for the 106 downregulated proteins in the Pi-limited *cnih5* roots. Enriched GO terms of (A) molecular function, (B) biological process, and (C) cellular component identified by ShinyGO were summarized using REVIGO to remove redundancy and cluster functionally related terms in *A. thaliana*. Each node represents a GO term, with color intensity reflecting Log_10_ (enrichment factor). The node size is proportional to the Log_10_ value of the number of gene annotations for each term in the background database. Lines between nodes represent semantic similarity, and line thickness indicates the degree of functional overlap between terms.

### Analysis of protein–protein interaction between *At*CNIH5 and its potential membrane cargoes

Considering that potential cargoes of *At*CNIH5 are downregulated integral MPs in *cnih5*, we narrowed the number of candidates to 31 (**Table 2**). This list is over-represented with three hypersensitive response-like lesion-inducing protein (HRLI)-like proteins (AT3G23175, AT3G23190, and AT4G14420), two *β*-ketoacyl-CoA synthases (*At*KCS2 and *At*KCS20), and two *At*PHT1 members (*At*PHT1;1 and *At*PHT1;3) (**Table 2**). Although *At*HRLI-like proteins have not been functionally characterized, the *N. benthamiana Nb*HRLI4 conferred salicylic acid-mediated resistance to turnip mosaic virus (TuMV) (Wu et al., 2021). *AtKCS* genes encode 3-ketoacyl-CoA synthases, which catalyze the rate-limiting condensation step of VLCFAs biosynthesis (Evenson and Post-Beittenmiller, 1995). *AtKCS2* and *AtKCS20* are highly expressed in the root endodermis and redundant in root suberin biosynthesis (Lee et al., 2009). Two alcohol-forming fatty acyl-CoA reductases, *At*FAR3/CER4 and *At*FAR5, are also downregulated and over-represented in the KEGG pathway of cutin, suberin, and wax biosynthesis (Fig. S4B, Table S2), suggesting that the biosynthesis of VLCFAs and their derivative extracellular aliphatic compounds is mis-regulated in the Pi-limited roots of *cnih5*. Nevertheless, *At*HRLI-like and *At*KCS proteins localize to the ER, so their downregulation may be a secondary effect of *At*CNIH5 impairment.

**Table 2.**
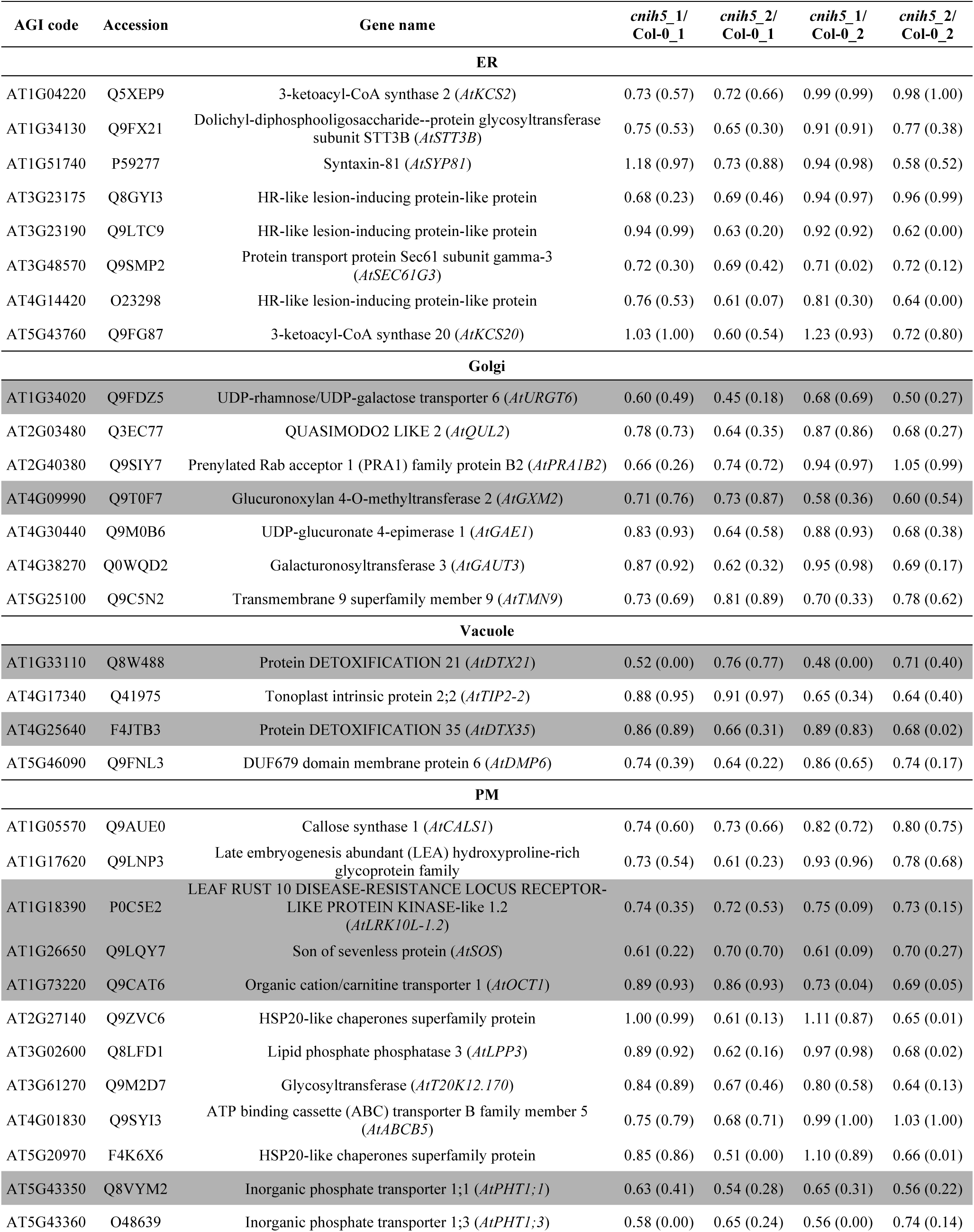
List of the 31 downregulated microsomal transmembrane proteins in Pi-limited *cnih5* roots. The normalized ratios compared the protein expression in Col-0_1, Col-0_2, *cnih5*_1 and *cnih5*_2 samples. The FDR adj. *p*-values are in parentheses. The candidates highlighted in gray are selected for testing protein–protein interaction in this study. Subcellular localization is predicted or annotated by SUBA5, UniprotKB or experimental data.

Next, we selected a subset of potential membrane cargoes based on the following two criteria (highlighted in gray in **Table 2**) to test their interaction with *At*CNIH5: (1) a TM protein located in post-ER compartments of the secretory pathway and (2) having four ratios ≤ 0.75 or at least two ratios ≤ 0.75 with at least one ratio showing FDR adj. *p*-value ≤ 0.05. We recently demonstrated that *At*CNIH5 interacts with *At*PHT1;1 and *At*PHT1;4 (Chiu et al., 2024). In this study, we further showed that *At*CNIH5 interacts with *At*PHT1;2, *At*PHT1;5, *At*PHT1;7, and *At*PHT1;9 in the yeast SUS (**Fig. 5A**), suggesting that *At*CNIH5 recognizes conserved motifs among *At*PHT1 members. In addition, we confirmed the interaction of *At*CNIH5 with the PM-localized receptor-like kinase *At*LRK10L-1.1, nucleotide exchange factor *At*SOS, and organic cation/carnitine transporter *At*OCT1, the Golgi-localized nucleotide sugar transporters *At*URGT6 and glucuronoxylan 4-O-methyltransferase *At*GXM2, and the tonoplast (TP)-localized detoxification efflux carriers *At*DTX21 and *At*DTX35 (**Fig. 5B)**. Similar results were also obtained when the *At*SOS, *At*OCT1, *At*URGT6, *At*DTX21, and *At*DTX35 expression constructs were generated with different fusion orientations (Fig. S5).

**Fig 5.**
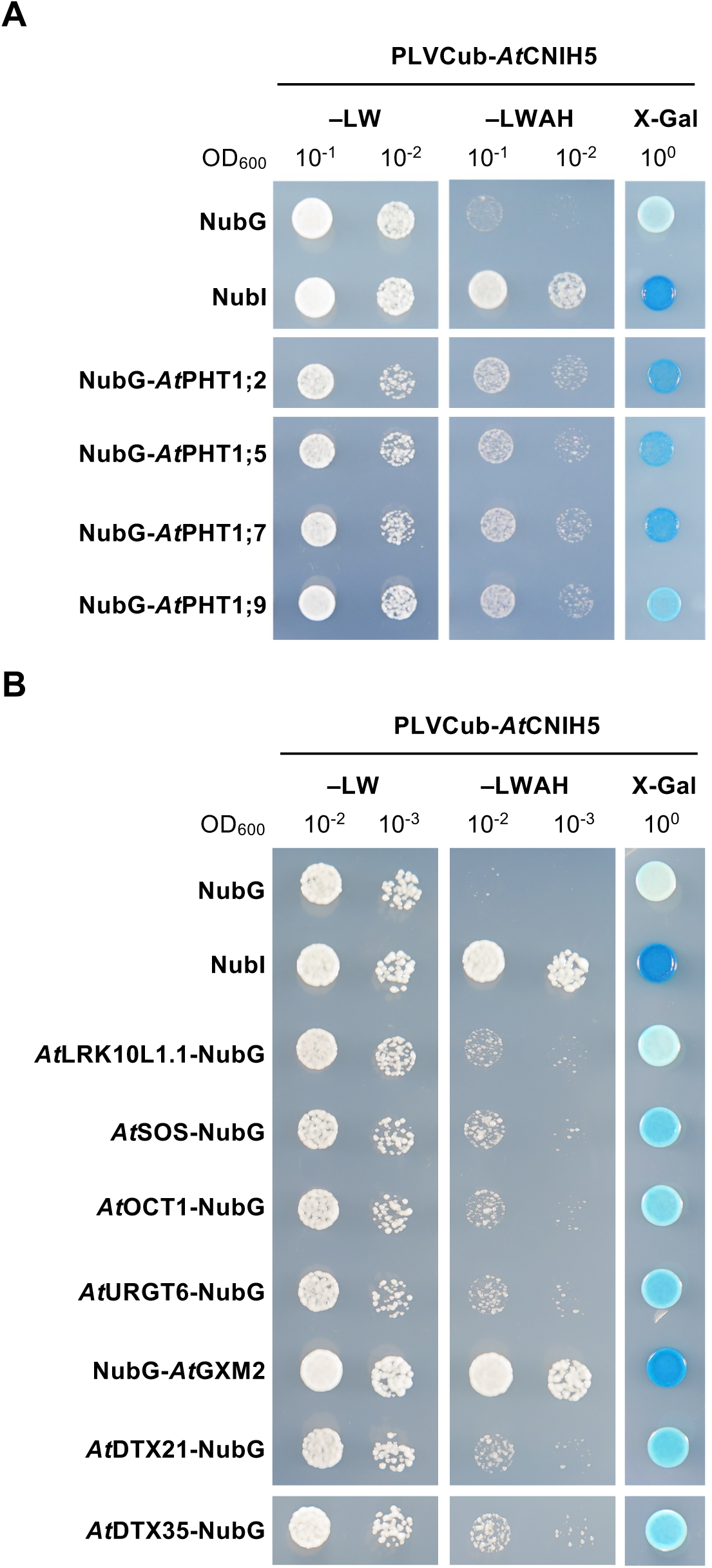
Analysis of the interaction of *At*CNIH5 with its potential ER cargoes by the yeast split-ubiquitin system. (A, B) Co-expression of PLVCub-*At*CNIH5 with NubG-*At*PHT1;2, NubG-*At*PHT1;5, NubG-*At*PHT1;7, and NubG-*At*PHT1;9 (A) or *At*LRK10L-1.1-NubG, *At*SOS-NubG, *At*OCT1-NubG, *At*URGT6-NubG, NubG*-At*GXM2, *At*DTX21-NubG, and *At*DTX35-NubG (B). The co-expression of PLVCub-*At*CNIH5 with NubI and NubG are used as positive and negative controls, respectively. Yeast transformants were grown on synthetic medium without leucine and tryptophan (–LW; the left panel) or on synthetic medium lacking leucine, tryptophan, adenine, and histidine with 500 μM methionine (–LWAH; middle panel) or on SC–LW containing 100 mg/L X-Gal (X-Gal; the right panel).

*AtOCT1* is a PSI gene (Liu et al., 2016) and its expression is restricted to the vasculature of the root, as well as at sites of lateral root formation (Lelandais-Brière et al., 2007), suggesting that the interaction of *At*CNIH5 and *At*OCT1 is likely to occur in the vascular tissues. Both *At*URGT6 and *At*GXM2 are cell wall-modifying enzymes. *At*URGT6 is involved in the biosynthesis of rhamnogalacturonan-I (RG-I), a component of pectin found in mucilage and various types of plant cell walls (Saez-Aguayo et al., 2020). *At*GXM2 is a methyltransferase that catalyzes 4-O-methylation of glucuronic acid (GlcA) side chains on the hemicellulose xylan (Lee *et al*., 2012; Urbanowicz *et al*., 2012; Yuan *et al*., 2014). Notably, *AtGXM2* is upregulated during Pi starvation (Liu et al., 2016) and is preferentially expressed in the root hair (Salazar-Henao and Schmidt, 2016). *At*DTX21 and *At*DTX35 belong to the multidrug and toxic compound extrusion (MATE)-type transporter superfamily. *At*DTX35 is also low Pi-responsive (Liu et al., 2016) and functions as a turgor-regulating chloride channel associated with root hair elongation (Zhang et al., 2017). Therefore, *At*CNIH5 may modulate cell wall elasticity and root hair growth, which is consistent with decreased root hair length in *cnih5* (Chiu et al., 2024).

To explore other potential membrane cargoes of *At*CNIH5 that may be missing in our iTRAQ-based proteomic data, we set out in parallel to find root PSI genes encoding transporters that are co-regulated with *AtCNIH5*. We queried the RNA-seq dataset from *Arabidopsis* seedlings subjected to Pi deprivation (P0: Pi-sufficiency; P1: one day of Pi deprivation; P3: three days of Pi deprivation) (Liu et al., 2016). Genes with expression levels less than 1 RPKM were excluded. We filtered root PSI genes that displayed an expression pattern similar to that of *AtCNIH5* based on (1) the log_2_FC ≥ 0.5 (P1/P0), (2) the log_2_FC ≥ 1 (P3/P0), and (3) the difference of log_2_FC ≥ 0.5 (P3/P0−P1/P0). Such cutoffs were applied to root PSI genes because the agriGO enrichment analysis (Tian et al., 2017) yielded a satisfactory list for further investigation, including nearly all *At*PHT1s and more than ten other genes encoding transporters (Table S4). Among the list, the PM-localized *At*OCT1, potassium transporter *At*HAK5, sulfate transporter *At*SULTR1;3, and the TP-localized Pi transporter *At*VPE1 were selected for testing interaction with *At*CNIH5 based on the tripartite split-GFP association in agro-infiltrated *N. benthamiana* leaves (Liu *et al*., 2018; Liu, 2021). We also included *At*SOS, *At*URGT6, *At*DTX21, and *At*DTX35 for testing, but excluded *At*LRK10L-1.1 and *At*GXM2 due to their similar membrane topology to *At*CNIH5 and the inability to find any agrobacteria transformants carrying the *At*LRK10L-1.1 construct. The S10-tagged *At*CNIH5 (S10-*At*CNIH5) and the S11-tagged fusion protein of interest correctly reached their subcellular destinations, except that *At*SOS-S11 and *At*SULTR1;3-S11 were retained in the ER (**Fig. 6A**). Compared to the recently reported positive control *At*PHT1;1-S11 (Chiu et al., 2024) and the negative control 3xHA-S11, S10-*At*CNIH5 also interacted with *At*OCT1-S11, *At*SOS-S11, *At*URGT6-S11, *At*DTX21-S11, and *At*DTX35-S11, yielding fluorescence signals as a result of GFP complementation; however, S10-*At*CNIH5 failed to interact with *At*HAK5-S11, *At*SULTR1;3-S11, and *At*VPE1-S11 (**Fig. 6B**). Taken together, our membrane proteomics successfully identified several potential cargoes of *At*CNIH5, including *At*PHT1s.

**Fig 6.**
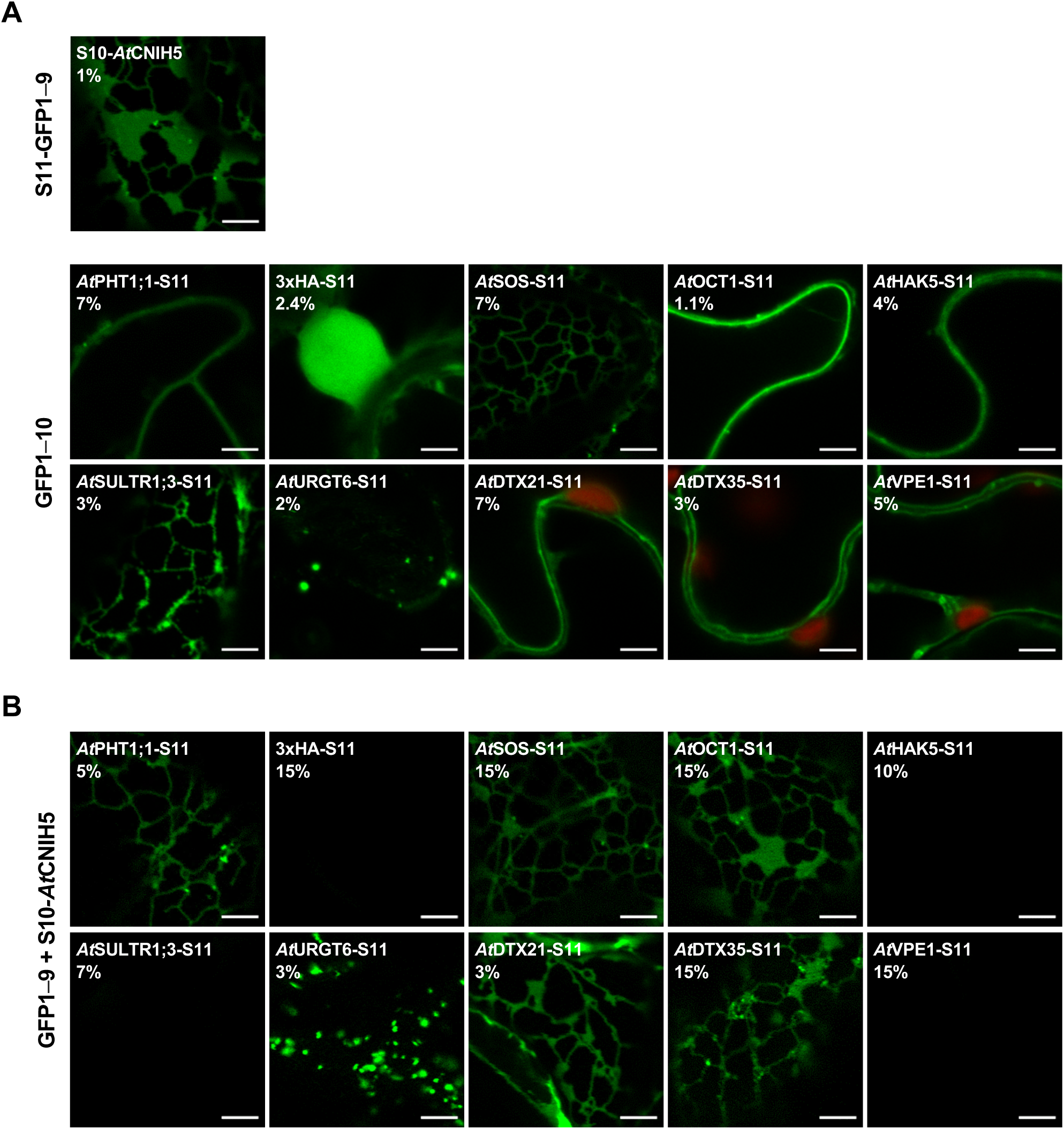
Analysis of the interaction of *At*CNIH5 with its potential ER cargoes by the tripartite split-GFP assay in *N. benthamiana* leaves. (A, B) Expression and subcellular localization (A) and protein–protein interaction analysis (B) of S10-*At*CNIH5 with *At*PHT1;1-S11, 3xHA-S11, *At*SOS-S11, *At*OCT1-S11, *At*HAK5-S11, *At*SULTR1;3-S11, *At*URGT6-S11, *At*DTX21-S11, *At*DTX35-S11, or *At*VPE1-S11. The co-expression of S10-*At*CNIH5 with *At*PHT1;1-S11 and 3xHA-S11 are used as positive and negative controls, respectively. The percentage of laser power used for acquisition is shown. Scale bars, 5 µm. Red signals show autofluorescence from chloroplasts.

### The C-terminal acidic residue of *At*CNIH5 is not required to interact with *At*PHT1;1

As *At*PHT1s were prominent cargoes of *At*CNIH5, we are interested in determining which motif of *At*CNIH5 interacts with *At*PHT1;1. On the basis of sequence similarity, we refer to the putative FLN motif in the second TMD of *At*CNIHs as the YLC motif and the putative cytosolic IFRTL motif as the IFNNL motif (Fig. S6). Notably, *At*CNIH5 has putative YLC and IFNNL motifs with several residues changed and a shorter C-terminal putative acidic motif with only one aspartic acid residue (Fig. S6). We designed a series of C-terminally truncated forms of *At*CNIH5. For an unknown reason, we failed to obtain the clone expressing *At*CNIH5 1–102. Therefore, we only tested the interaction of *At*PHT1;1 with *At*CNIH5 1–26, *At*CNIH5 1–49, *At*CNIH5 1–92, and *At*CNIH5 1–132 in the yeast SUS (**Fig. 7A**). *At*CNIH5 1–26, which contains only the first TMD, is sufficient to interact with *At*PHT1;1, while *At*CNIH5 1–132, which has the last three amino acids deleted, including the C-terminal aspartic acid, still interacts with *At*PHT1;1 (**Fig. 7B**). We also examined whether the truncated variants *At*CNIH5 1–26 and *At*CNIH5 1–132 interact with *At*PHT1;1 in plant cells. Both the *At*CNIH5 truncated variants localized to the ER (**Fig. 7C**) and interacted with *At*PHT1;1 (**Fig. 7D**). Interestingly, *At*CNIH5 1–132 also interacted with *At*DTX21 but not with *At*OCT1 (**Fig. 7C**). Therefore, the C-terminal acidic residue of *At*CNIH5 is required for interaction with *At*OCT1 but not with *At*PHT1;1 or *At*DTX21. As such, *At*CNIH5 interacts with *At*PHT1;1, and likely the other *At*PHT1s, through a distinct mechanism that differs from those used by the fungal and rice CNIHs for their cognate cargoes.

**Fig 7.**
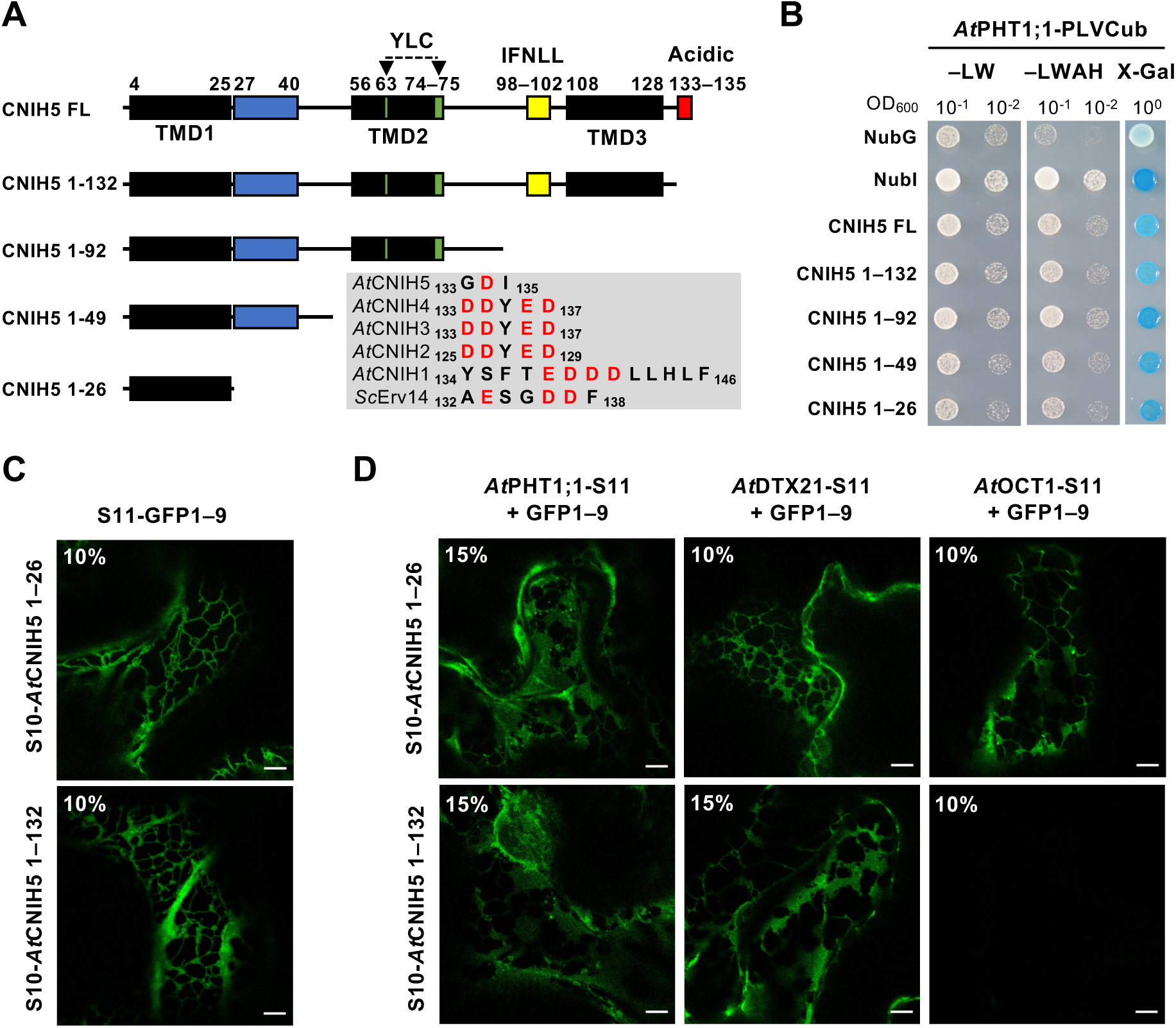
Analysis of the interaction of *At*PHT1;1 with the truncated *At*CNIH5 variants. (A) Schematic diagram of full-length (FL) and truncated *At*CNIH5 variants. The lengths of the truncated variants are indicated by the number of residues. The transmembrane domains and the putative cornichon, YLC, IFNLL, and acidic motifs are shown. The black box, the transmembrane domain (TMD); the blue box, the cornichon motif; the green box, the YLC motif; the yellow box, the IFNLL motif. The red box, the acidic motif. The gray box shows the sequence alignment of the putative acidic motif of *Sc*Erv14 and *At*CNIHs. Noted that *At*CNIH5 has a shorter C-terminal tail carrying only one acidic residue D134. (B) Co-expression of *At*PHT1;1-PLVCub with NubG-*At*CNIH5, NubG-*At*CNIH5 1–26, NubG-*At*CNIH5 1–49, NubG-*At*CNIH5 1–92, and NubG-*At*CNIH5 1–132 in yeast split ubiquitin assay. The co-expression of *At*PHT1;1-PLVCub with NubI and NubG are used as positive and negative controls, respectively. Yeast transformants were grown on synthetic medium without leucine and tryptophan (–LW; the left panel) or on synthetic medium lacking leucine, tryptophan, adenine, and histidine with 500 μM methionine (–LWAH; middle panel) or on SC–LW containing 100 mg/L X-Gal (X-Gal; the right panel). (C) Expression and subcellular localization of S10-*At*CNIH5 1–26 and S10-*At*CNIH5 1–132 in *N. benthamiana* leaves. (D) Protein–protein interaction analysis of *At*PHT1;1-S11, *At*DTX21-S11, or *At*OCT1-S11 with S10-*At*CNIH5 1–26 or S10-*At*CNIH5 1–132 in *N. benthamiana* leaves. The percentage of laser power used for acquisition is shown. Scale bars, 5 µm.

### Increasing *in-situ At*CNIH5 expression/activity enhances plant growth

Since loss of *At*CNIH5 leads to decreased abundance of *At*PHT1s and other membrane cargoes (**Table 2**), it was imperative to assess whether overexpression of *At*CNIH5 can improve plant fitness. We found that ectopic expression of RFP-*At*CNIH5 driven by the cauliflower mosaic virus (CaMV) 35S promoter resulted in distorted root morphology (Fig. S7). This aberrant structure was likely caused by ER deformation, a phenomenon associated with Erv14 overexpression in budding yeast (Lagunas-Gomez et al., 2023). Intriguingly, we recently showed that several transgenic lines expressing the genomic GFP-*AtCNIH5* fragment (*AtCNIH5pro*: GFP-*AtCNIH5*) in the *cnih5* background restored the abundance of *At*PHT1;1/2/3 to levels comparable to or higher than WT during Pi deficiency (Chiu et al., 2024). Under Pi sufficiency, these transgenic lines grew larger and showed WT-like shoot Pi content *per plant,* but had lower shoot Pi levels (Chiu et al., 2024). Therefore, we systematically examined the growth of these lines under different Pi regimes. Under Pi sufficiency and modest Pi deficiency—either three days of Pi deprivation (0 µM Pi) or eleven days of suboptimal Pi supply (50 µM Pi), these plants exhibited higher shoot and/or root fresh weight than the WT and *cnih5* plants (**Fig. 8**). In contrast, under more severe Pi deficiency—eleven days of Pi deprivation (0 µM Pi), none of these lines had a larger shoot size than WT (**Fig. 8**). Although we could detect GFP-*AtCNIH5* proteins in the Pi-limited root of *AtCNIH5pro*: GFP-*AtCNIH5/cnih5* plants, it is technically difficult to discern whether GFP-*AtCNIH5* proteins were more abundant than the endogenous *At*CNIH5 in WT (Fig. S8). Nonetheless, the higher biomass of these transgenic plants is likely due to increased expression or activity of GFP-*AtCNIH5*, along with additional membrane cargoes beyond *At*PHT1s, leading to more efficient assimilation of acquired Pi. We inferred that a subtle increase in *in-situ* rather than ectopic expression/activity of GFP-*AtCNIH5* suffices to activate downstream effectors of *At*CNIH5, thereby boosting plant growth.

**Fig 8.**
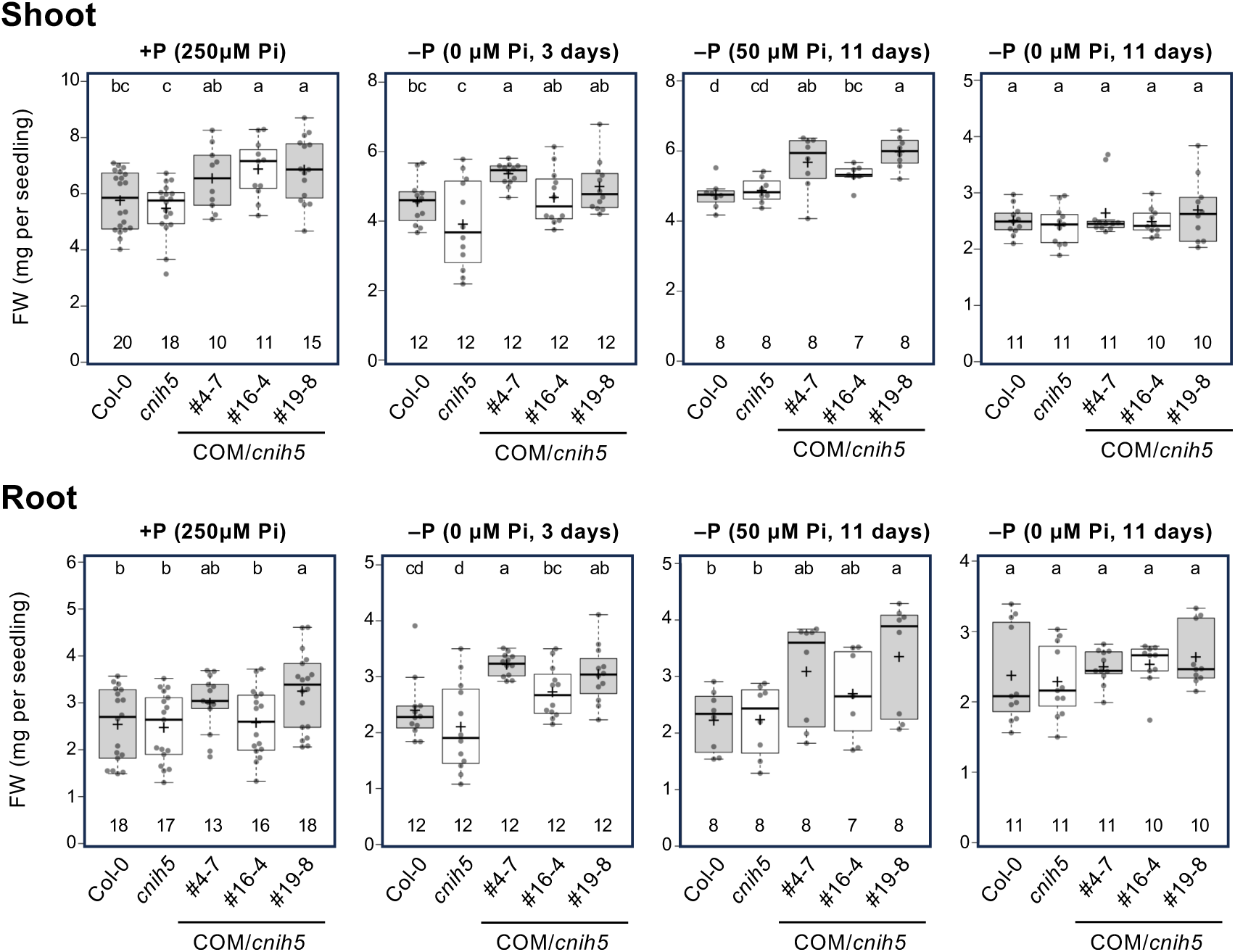
Increasing *in-situ At*CNIH5 expression/activity boosts plant growth. The shoot and root fresh weight (FW) of *Arabidopsis* 11-day-old WT, *cnih5* and *AtCNIH5pro*: GFP-*AtCNIH5/cnih5* complementation (COM) lines under +P (250 µM KH_2_PO_4_), modest –P (0 µM KH_2_PO_4_, three days of starvation or 50 µM KH_2_PO_4_, eleven days of starvation), and severe –P (0 µM KH_2_PO_4_, eleven days of starvation) conditions. Data are visualized using BoxPlotR (Spitzer *et al*., 2014). Center lines show the medians; the central plus signs (+) show the means; box limits indicate the 25th and 75th percentiles; whiskers extend to the 5th and 95th percentiles, and data points are plotted as dots. Data significantly different from the other groups are indicated by different lowercase letters (n, pools of seedings from independent experiments; One-Way ANOVA, Duncan’s test).

## Discussion

CNIHs are the eukaryotic conserved cargo receptors that cycle between the ER and the early Golgi to select membrane proteins for incorporation into COPII vesicles. In this study, we aimed to identify potential membrane cargoes of low Pi-induced *At*CNIH5 through iTRAQ-based proteomic analysis, while also providing the first comprehensive membrane proteome for *Arabidopsis* roots subjected to Pi deprivation (Zhou et al., 2022). Given that our iTRAQ-based proteomic analysis did not include Pi-sufficient WT and *cnih5* samples, we sought to evaluate whether the identified potential membrane cargoes of *At*CNIH5 are PSI proteins. To address this, we examined the overlap between the proteins we identified in the Pi-limited roots and the root PSI genes in the RNA-seq dataset (P3/P0, FC > 1.5; Liu et al., 2016). We found minimal overlap, at only 5.8%, between the 4,317 identified proteins and the root PSI genes (Table S1). This finding aligns with the previous study, which indicated a weak correlation between transcript and protein changes following Pi deficiency in *Arabidopsis* roots (Lan et al., 2012). Still, of our identified integral MPs in the Pi-limited roots, 7.8% (4 out of 51) of the upregulated and 32.3% (10 out of 31) of the downregulated proteins in *cnih5* were classified as root PSI genes (Table S5). These results indicated that, compared with PSI genes within the 4,317 identified proteins, PSI genes are enriched 5.6-fold among the downregulated integral MPs in Pi-limited *cnih5* root (Table S5). Therefore, the *At*CNIH5-dependent membrane cargoes we identified are mainly associated with Pi starvation.

When expressed in the yeast heterologous expression system, *At*CNIH5 interacted not only with the two major Pi transporters, *At*PHT1;1 and *At*PHT1;4 (Chiu et al., 2024), but also with *At*PHT1;2, *At*PHT1;5, *At*PHT1;7, and *At*PHT1;9 (**Fig. 5A**), supporting that *At*PHT1s are the prominent membrane cargoes of *At*CNIH5. This was not surprising because the amino acid sequences of *At*PHT1s share over 47.4% identity. It has been shown that the yeast and rice CNIHs used the C-terminal acidic motif to interact with their cognate cargoes (Rosas-Santiago et al., 2017). However, a recent study revealed that the cargo binding motif of the pumpkin *Cm*CNIH1 may reside in the N-terminal rather than the C-terminal region (Wei et al., 2023). This agreed with our results for the interaction between *At*CNIH5 and *At*PHT1;1 (**Fig. 7B**), indicating that the first TMD of *At*CNIH5 contains a motif that interacts with *At*PHT1s.

Besides *At*PHT1s, we identified other PM-localized membrane proteins that were downregulated in Pi-limited *cnih5* roots. *At*LRK10L-1.2 is involved in ABA signaling (Lim et al., 2015). The animal Son of Sevenless (SOS) proteins encode guanine nucleotide exchange factors that act on the Ras subfamily of small GTPases (Bowtell et al., 1992). However, the physiological role of SOS in plants remains to be studied. The PSI *At*OCT1 was also identified as a cargo of *At*CNIH5. We further demonstrated that the C-terminal motif of *At*CNIH5 is required for the interaction between *At*OCT1 and *At*CNIH5 (**Fig. 7D**). Notably, the *oct1* mutant exhibited enhanced root branching (Lelandais-Brière et al., 2007), phenocopying the higher lateral root density of *cnih5* (Chiu et al., 2024). Although the heterologous expression of *At*OCT1 in yeast could transport carnitine, it remains to be determined which substrate *At*OCT1 transports that could negatively affect lateral root development.

Previous analyses of root-specific co-expression have indicated that early PSR genes are involved in cell wall remodeling, highlighting the importance of this process in response to Pi deficiency (Liao *et al*., 2011; Lin *et al*., 2011). Various Golgi-localized proteins involved in plant cell wall modification, such as *At*QUL2 (AT2G03480) (Mouille et al., 2007), *At*GAE1 (AT4G30440) (Mølhøj et al., 2004), *At*GAUT3 (AT4G38270) (Sterling et al., 2006), and *At*TMN9 (AT5G25100) (Hiroguchi et al., 2021), were remarkably downregulated in the Pi-limited *cnih5* roots (**Table 2**). The cytosolic UDP-glucose pyrophosphorylase 2 (UGP2, AT5G17310), which is implicated in the synthesis of cell wall and callose deposition, was also downregulated in *cnih5* across the four ratios (**Fig. 3D**). In contrast, the Golgi-localized hydroxyproline-O-galactosyltransferase *At*GALT9 (AT1G53290), which is involved in pectin metabolism (Zhang et al., 2021), was the most highly upregulated protein in Pi-limited *cnih5* roots (**Fig. 3D**). We speculate that *At*CNIH5 may play a critical role in Pi starvation-triggered cell wall composition changes. Nevertheless, the difference in cell wall structure between WT and *cnih5* remains to be investigated.

The TP-localized *At*DTX21 and *At*DTX35 were identified as the potential cargoes of *At*CNIH5 (**Fig. 5B**, **6B**). While *AtDTX35* is known as a PSI gene, *AtDTX21* is co-induced with the type III effectors secreted by *Pseudomonas syringae* and mediates the detoxification of chlorinated herbicide atrazine (de Torres-Zabala *et al*., 2007; Ramel *et al*., 2007). Further studies are needed to elucidate the physiological role of *At*DTX21 and *At*DTX35 in Pi starvation responses. *At*CNIH5 failed to interact with other manifest PSI transporters such as *At*HAK5, *At*SULTR1;3, and *At*VPE1 (**Fig**. **5B**). In fact, *At*VPE1 and its paralog *At*VPE3 were identified in our proteomic analysis, while their abundance did not differ between WT and *cnih5* (Table S1). This suggests that *At*CNIH5 recognizes only specific PSI transporters via a mechanism yet to be elucidated.

In conclusion, an Azo-solubilized MME approach in conjunction with quantitative proteomics was used to reveal the downregulation of *At*PHT1s and other membrane proteins associated with Pi starvation in *cnih5* roots (**Fig. 9**). Our analyses validated the interaction between *At*CNIH5 and several membrane proteins at post-ER compartments, including *At*PHT1s, *At*OCT1, *At*URGT6, *At*DTX21, and *At*DTX35 transporters. We also showed that the protein level of *At*CNIH5 increases during Pi starvation as well as in the Pi-replete root of *pho2*, as a result of accommodating the increased demand for ER export of *At*PHT1s. Moreover, introducing the genomic GFP-*AtCNIH5* fragment enhanced plant growth across a wide range of Pi supply levels. Altogether, we propose that *At*CNIH5 acts as a Pi deficiency-inducible hub, facilitating the ER-to-Golgi trafficking of several membrane proteins through a mechanism beyond the putative C-terminal acidic motif, thereby improving plant fitness under suboptimal Pi availability.

**Fig. 9.**
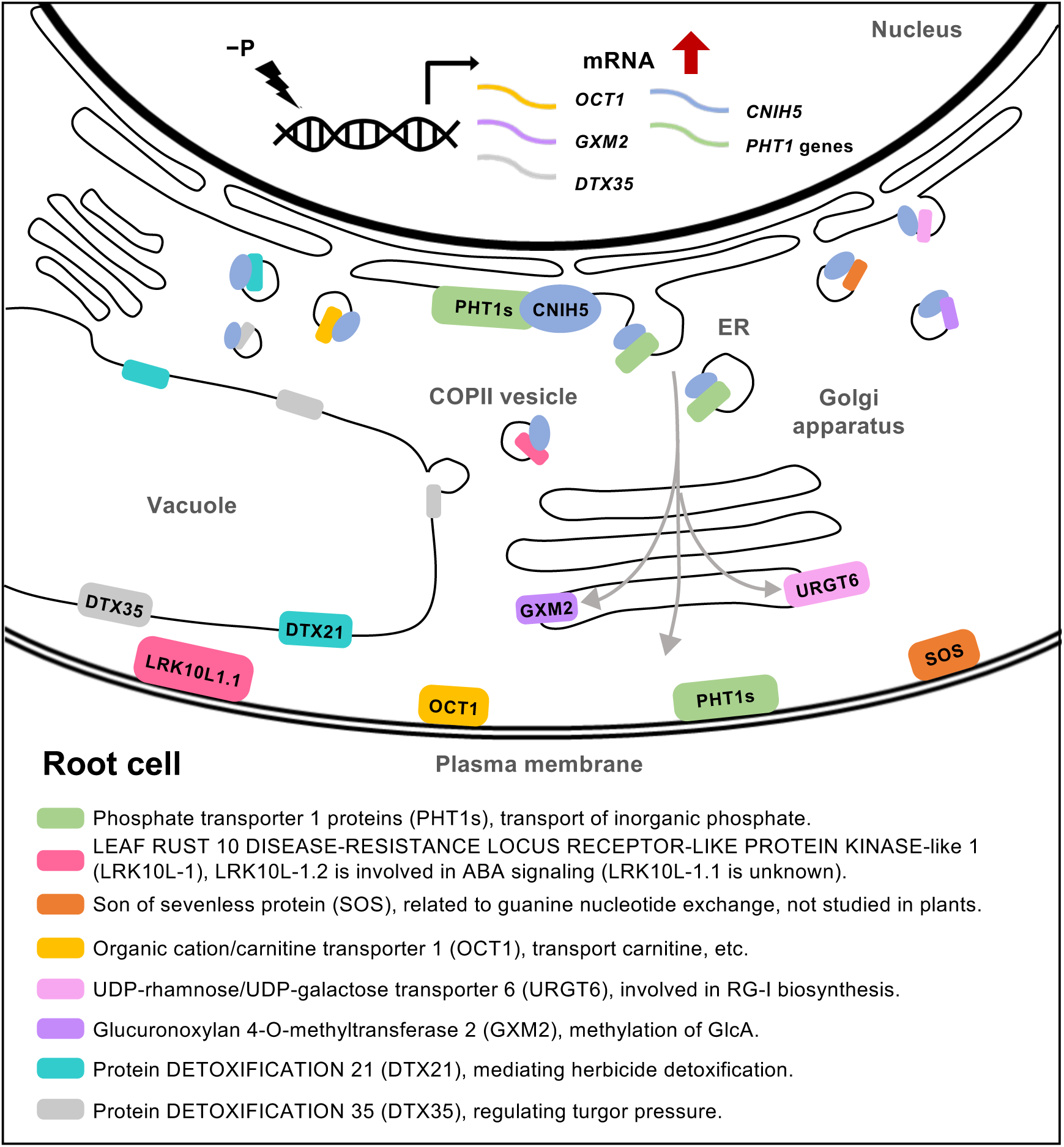
*At*CNIH5-mediated ER-to-Golgi transport of membrane proteins associated with Pi starvation responses in *Arabidopsis* root. Under Pi starvation (−P), the transcripts of *CNIH5*, *PHT1* genes, *OCT1*, *GXM2*, and *DTX35* increase in *Arabidopsis* root cells. As an ER cargo receptor, CNIH5 assists the ER export of the newly synthesized PHT1s, LRK10L-1.1, SOS, OCT1, URGT6, GXM2, DTX21, and DTX35, and their loading into COPII vesicles, thereby directing them to their corresponding subcellular destinations.

## Materials and Methods

### Plant Material and Growth Conditions

Seeds of the *A. thaliana* wild-type (Columbia, Col-0) and *cnih5* (SAIL_256_B03) mutants used in this study were obtained from the Arabidopsis Biological Resource Center. Seeds were surface-sterilized and germinated on agar plates with one-half modified Hoagland’s solution containing 1% sucrose and 0.7% Bacto agar and grown in the growth chamber at 22 °C with a 16-h light/8-h dark cycle. Pi-sufficient (+P; 250 μM Pi) or Pi-deficient (−P; 0 μM Pi) medium containing 1.2% agar was used for vertically grown plants.

### Tissue Homogenization and Microsomal Protein Preparation

The roots of 11-day-old seedlings with five days of Pi starvation were harvested. Approximately 100 mg of root tissue was collected for each sample and stored at −80 °C for subsequent homogenization. The frozen samples were homogenized by TissueLyser II (QIAGEN). The method for microsomal protein extraction was modified from a previous study (Abas and Luschnig, 2010), and all steps were performed on ice or 4 ℃.

Briefly, each sample was resuspended in 100 μl of extraction buffer (EB) containing 50 mM of hydroxyethyl piperazine ethane sulfonic acid (HEPES; Melford, pH 7.5), 25% (w/w 0.81 M) sucrose, 5% (v/v) glycerol, 10 mM of ethylenediaminetetraacetic acid (EDTA; Amresco, pH 8), 5 mM of KCl, 1 mM of dithiothreitol (DTT; Sigma), and 2× protease inhibitor cocktail (P9599, Sigma). An equivalent volume (100 μl) of 5% polyvinylpolypyrrolidone (PVPP; Sigma) suspension solution was used for each sample. The 5% PVPP was centrifuged at 2,000×g for 1 min at room temperature (RT). The precipitate was activated for 2 hours in an equivalent volume of activated solution containing 100 mM HEPES, 40% (w/w, 1.37 M) sucrose, 20 mM EDTA, and 10 mM KCl. The activated PVPP was centrifuged at 5,000×g for 1 min at RT, and the supernatant was removed. The activated PVPP pellet was mixed with the sample suspension and stood for 5 min, followed by centrifugation thrice. The first centrifugation was processed at 600×g for 3 min. The pellet was then re-extracted with 50 μl EB and centrifuged at 600×g for 3 min. The pellet was then re-extracted with 35 μl EB and centrifuged at 2,000×g for 30 s. After three times centrifugation, all the supernatants were combined and centrifuged again at 600×g for 3 min. The collected supernatant was diluted with an equivalent volume of ddH_2_O to reach a final solution concentration containing 12–13% sucrose and centrifuged at 21,000×g for 90 min. The pellet was resuspended in washing buffer containing 10 mM HEPES, 5 mM EDTA, 1 mM phenylmethylsulfonyl fluoride (PMSF, Sigma), and 1× protease inhibitor cocktail and centrifuged at 21,000×g for 45 min. Finally, the pellet was stored at −80 ℃ for further analysis.

### Protein Solubilization and In-Solution Digestion

Microsomal protein pellets were dissolved in 1% sodium 4-hexylphenylazosulfonate (Na-Azo, Excenen Pharmatech) in 100 mM triethylammonium bicarbonate (TEABC, Sigma) containing 2× protease inhibitor cocktail and 1 mM PMSF. The protein yield was determined by BCA assay (Pierce). For each sample, an equal amount of protein was reduced with 5 mM tris-(2-carboxyethyl)-phosphine (TCEP, Sigma) at 60 °C for 30 min, followed by cysteine-blocking with 10 mM methyl methanethiosulfonate (MMTS, Sigma) at 25 °C for 30 min. Samples were digested with sequencing-grade modified porcine trypsin (Promega) at 37 °C with shaking for 1 hour and then illuminated by 6W UV light 302 nm (UVP UVM-57, Analytik Jena) at 4 °C for 40 min. The digested proteins were then aliquot and stored at −20 °C until use.

### Isobaric Tag for Relative and Absolute Quantitation (iTRAQ) Labeling and Peptide Separation

After trypsin digestion, 0.5 µg of peptides was subjected to 1D–LC–MS/MS without pre-fractionations to identify and quantify proteins as a quality control study. On the other hand, 10 µg of peptides was labeled using an iTRAQ Reagent-4 plex Multiplex Kit according to the manufacturer’s instructions (Applied Biosystems) for 1 hour at RT. Two WT replicates were labeled with iTRAQ-114 and -115, and two *cnih5* replicates were labeled with iTRAQ-116 and -117, pooled and desalted by homemade C18-microcolumn (SOURCE 15RPC, GE Healthcare). The eluted peptides were dried by vacuum centrifugation. The desalted peptides were loaded onto a homemade strong cation exchange (SCX) column (Luna SCX 5μm, 0.5 × 120 mm) at a 2 μl/min flow rate for 60 min. The peptides were then fractionated to 24 fractions by eluting with 0 to 100% HPLC buffer B (1 M ammonium nitrate/25% acetonitrile/0.1% formic acid) using on-line two-dimensional high-performance liquid chromatography (2D-HPLC; Dionex Ultimate 3,000, Thermo Fisher). Each SCX fraction was diluted in-line before trapping onto a reverse-phase (RP) column (Zorbax 300SB-C18, 0.3 x 5 mm; Agilent Technologies). The peptides were then separated on a column (Waters BEH 1.7 μm, 100 μm I.D. × 10 cm with a 15 μm tip) using a multi-step gradient of HPLC buffer C (99.9% acetonitrile/0.1% formic acid) for 60 min with a flow rate of 0.3 μl/min.

### Liquid Chromatography-Tandem Mass Spectrometry (LC-MS/MS) Analysis

The LC apparatus was coupled with a 2D linear ion trap mass spectrometer (LTQ-Orbitrap ELITE; Thermo Fisher) operated using Xcalibur 2.2 software (Thermo Fisher). The full-scan MS was performed in the Orbitrap over a range of 400 to 2,000 Da and a resolution of 120,000 at m/z 400. For proteome analysis, the 10 data-dependent MS/MS scan events were followed by one MS scan for the 10 most abundant precursor ions in the preview MS scan. The m/z values selected for MS/MS were dynamically excluded for 80 sec with a relative mass window of 1 Da. The electrospray voltage was set to 1.8 kV, and the capillary temperature was set to 200 °C. MS and MS2 automatic gain control were set at 1,000 ms (full scan) and 300 ms (MS2 for HCD), or 3 × 10^6^ ions (full scan) and 3 × 10^4^ ions (MS2 for HCD) for maximum accumulated time or ions.

### LC-MS Data Processing, Protein Identification and Quantification

Proteome Discoverer software (version 2.3, Thermo Fisher Scientific) was used for data analysis, incorporating the reporter ions quantifier node for iTRAQ quantification. The MS/MS spectra were searched against the SwissProt database using the Mascot search engine (Matrix Science, version 2.5). For peptide identification, a 10 ppm mass tolerance was allowed for intact peptide masses and 0.2 Da for HCD fragment ions, with allowance for two missed cleavages made from trypsin digestion. Variable modifications included oxidized methionine, acetylation (protein N-terminal), and iTRAQ-4plex (peptide N-terminal, K); methylthiolation (C) was considered a static modification. Peptide-spectrum matches (PSMs) were filtered based on high confidence and Mascot search engine rank 1 of peptide identification, ensuring an overall false discovery rate (FDR) below 0.01. Proteins with only a single peptide hit were removed. Quantitative protein data were exported from Proteome Discoverer using Integration Methods of the Most Confident Centroid with a 20 ppm tolerance. The data sets were analyzed by cross-comparison between four samples (ratio 1, 116:114; ratio 2, 117:114; ratio 3, 116:115, and ratio 4, 117:115). Protein ratios in each comparison group were normalized using the median ratio of each comparison group. Volcano plots of differentially expressed proteins were generated using the ggplot2 (Wickham et al., 2016) and ggrepel (Slowikowski et al., 2024) package in R version 4.3.1 (R Core Team, 2021). The code was written using RStudio version 2023.06.2 (RStudio Team, 2020). A 1.25-fold cutoff was applied to define differentially expressed proteins in each comparison group. The mass spectrometry proteomics data have been deposited to the ProteomeXchange Consortium via the PRIDE partner repository (Deutsch et al., 2022; Perez-Riverol et al., 2024) with the dataset identifier PXD062137 and 10.6019/PXD062137.

### Protein Annotation and Interpretation

Transmembrane-related information was obtained through analysis using DeepTMHMM (https://dtu.biolib.com/DeepTMHMM) (Hallgren et al., 2022). Subcellular structure localization was analyzed by SUBA5 (https://suba.live/) (Hooper et al., 2016). For functional annotation of proteins, Gene Ontology (GO) and Kyoto Encyclopedia of Genes and Genomes (KEGG) enrichment analyses were determined by ShinyGO 0.82 (http://bioinformatics.sdstate.edu/go/) (Ge et al., 2019), in which *p*-values were computed using the hypergeometric test and multiple test correction was performed using the Benjamini–Hochberg method based on an FDR cut-off of 0.05. Only pathways with a minimum of 10 genes were selected. Similar pathways that share 95% of their genes and 50% of the words in their names were eliminated according to the setting of ShinyGO 0.82. Dot plots of GO enrichment analysis were generated using the ggplot2 (Wickham et al., 2016) and stringr package (Wickham, 2023).

### Microsomal protein extraction for immunoblot analysis

The MPs pellets were solubilized by 0.5% Azo in 25mM aqueous ammonium bicarbonate, pH 7.0 (Sigma Aldrich). Alternatively, the low-speed pellet (LSP) was prepared as previously described (Yoshimoto et al., 2004), and microsomal protein was isolated using the Minute Plant Microsomal Membrane Extraction Kit (Invent, MM-018) according to the manual instructions. A 5–30 μg of MPs from each sample was loaded onto 4–12% Q-PAGE™ Bis-Tris Precast Gel (SMOBIO) or NuPAGE 4–12% Bis-Tris Gels (Thermo Fisher Scientific). Gel was transferred to polyvinylidene difluoride membranes (Immobilon-P Membrane or Immobilon®-PSQ Membrane; Merck). Membrane was blocked with 1–2% BSA in 1× PBS solution with 0.2% Tween 20 (PBST, pH 7.2) at RT for 1 h, and rinsed two times with distilled water followed by hybridization with primary antibodies of *At*PHF1 (Huang et al., 2013), *At*PHO1 (Liu et al., 2012), *At*NRT2.1 (AS12 2612; Agrisera), and *At*CNIH5 at RT for 2 h or at 4 °C overnight in blocking solution. Polyclonal rabbit *At*CNIH5 antibody was raised against the region corresponding to the amino acid residues 96 to 114 (TELYNTNKWEQKKRVYKIG) by LTK BioLaboratories, Taiwan, and was affinity purified. The membrane was washed four times with 1× PBST for 5 min followed by hybridization with the horseradish peroxidase–conjugated secondary antibody (GTX213110-01 or GTX213111-01; GeneTex) in blocking solution at RT for 1 h. After four washes in 1× PBST for 5 min and a rinse with distilled water, chemiluminescent substrates (WesternBright ECL or Sirius; Advansta) were applied for signal detection.

### Construct design and yeast split-ubiquitin assay

For constructing the yeast expression clones, the coding sequence (CDS) of the gene of interest (GOI), including *At*LRK10L-1.1, *At*GXM2, *At*SOS*, At*DTX21*, At*DTX35*, At*URGT6, *At*OCT1, and the truncated *At*CNIH5 variants, was amplified by PCR, cloned into the pCR8/GW/TOPO vectors and sequenced. The pCR8/GW/TOPO construct containing the CDS of *AtPHT1;1* was cloned as previously reported (Huang et al., 2013). The entry clones were recombined into NX32_GW (ABRC CD3-1737) or XN22_GW (ABRC CD3-1735) through LR reactions, resulting in N or C-terminally fused Nub fragments or into MetYC_GW (ABRC CD3-1740) through LR reactions, resulting in *At*PHT1;1-PLVCub. The PLVCub-*At*CNIH5 construct was generated as recently reported (Chiu et al., 2024). The Nub and PLVCub expression clones were sequentially transformed into the yeast THY.AP4 strain (ABRC CD3-808; (Obrdlik et al., 2004). Fresh yeast colonies picked from yeast extract peptone dextrose adenine (YPDA) medium or synthetic complete (SC) medium without leucine (SC–L) were inoculated in the corresponding liquid medium and incubated overnight at 30 ℃ with shaking at 200 rpm. Overnight yeast culture was then diluted to OD_600_ = 0.2 and grew for another 4–6 h until OD_600_ reached 0.4–0.6. Yeast pellet was collected by centrifugation at 2,000×g at 4 ℃ for 10 min, washed with ddH_2_O, and resuspended in Tris-EDTA/lithium acetate solution (TE/LiAc, 10 mM Tris-HCl [pH 8.0], 1 mM EDTA, and 0.1 M lithium acetate). Yeast transformants carrying the PLVCub*-At*CNIH5 or *At*PHT1;1-PLVCub were selected and maintained on SC–L for sequential transformation with NubI (positive control), NubG (negative control), and GOI-NubG or NubG-GOI. Yeast transformants carrying the Nub and the PLVCub constructs were selected and maintained on SC medium without leucine and tryptophan (SC–LW). For the protein–protein interaction testing, yeast cells co-expressing PLVCub*-At*CNIH5 or *At*PHT1;1-CubPLV with GOI-NubG or NubG*-*GOI were grown on synthetic media lacking leucine, tryptophan, adenine, and histidine (SC–LWAH) with 500 μM methionine or on SC–LW containing 100 mg/L 5-bromo-4-chloro-3-indolyl-*β*-d-galactopyranoside (X-Gal) at 30 ℃ for 24–36 h. Primer sequences used are listed in Table S6.

### Construct design for split-GFP assays in *N. benthamiana* leaves and *Arabidopsis* transgenic plants

The *UBQ10:sXVE:* S10-*At*CNIH5 clone was obtained as recently reported (Chiu et al., 2024), and *UBQ10:sXVE:* S10-*At*CNIH5 1–26 and *UBQ10:sXVE:* S10-*At*CNIH5 1–132 were obtained through subcloning. For the transient expression of *At*PHT1;1-S11, *At*SOS-S11, *At*DTX21-S11, *At*DTX35-S11, *At*URGT6-S11, *At*OCT1-S11, *At*HAK5-S11, *At*VPE1-S11, and *At*SULTR1;3-S11, the CDS or the corresponding fragment was amplified by PCR and subcloned into *UBQ10:sXVE*: (MCS)-S11 (Addgene plasmid # 108179). For subcellular localization analysis of the protein of interest, either *UBQ10:sXVE:* S11-GFP1–9 (Addgene plasmid # 125665) or *UBQ10:sXVE:* GFP1–10 (Addgene plasmid # 125663) was co-expressed. For the tripartite split-GFP reconstitution, the constructs encoding S10-tagged and S11-tagged protein fusions were co-expressed in the presence of *UBQ10:sXVE*: GFP1–9 (Addgene plasmid # 108187). Co-expression with 3xHA-S11 (Addgene plasmid # 108238) was used as a negative control for protein–protein interaction. The brightness of the images for the protein–protein interaction was adjusted by a 20% increase, and the contrast adjusted by a 20% decrease. For the genomic GFP-*AtCNIH5* complemented plants (*AtCNIH5pro*: GFP-*AtCNIH5/cnih5*), the construct design was recently described (Chiu et al., 2024). The plasmid encoding *35S*: RFP-*At*CNIH5 was obtained as previously reported (Wudick *et al*., 2018). Primer sequences used are listed in Table S6.

### Agrobacterium tumefaciens-mediated infiltration of N. benthamiana leaves

*Agrobacterium tumefaciens* strain EHA105 harboring the expression constructs or p19 was cultured in Luria–Bertani (LB) medium with the appropriate antibiotics [gentamicin (50 μg ml^−1^), rifampicin (5 μg ml^−1^) or kanamycin (50 μg ml^−1^)]. Cells were resuspended in the infiltration medium containing 10 mM MgCl_2_, 10 mM MES (pH 7.5), and 150 μM acetosyringone, and co-infiltrated to the true leaves of 3–5-week-old tobacco plants. For the *β*-estradiol-inducible split-GFP system, 36.7 μM *β*-estradiol was applied as previously described (Liu *et al*., 2018; Liu, 2021). Infiltrated leaves were collected 2 days post-infiltration (dpi) for confocal imaging.

### Confocal microscopy

Confocal microscopy images were acquired using Zeiss LSM 800 (Zeiss) with objectives Plan-Apochromat 10×/0.45 M27, 20×/0.8 M27, and 40×/1.3 Oil DIC M27 in multi-track mode with line switching and averaging of two readings. Excitation/emission wavelengths were 488 nm/410–546 nm for GFP, and 561 nm/656–617 nm for RFP, and 640 nm/656–700 nm for chlorophyll autofluorescence.

### Chemical treatments

Acetosyringone (150 mM stock in DMSO; D134406; Sigma-Aldrich) was diluted to a final concentration of 150 µM in ddH_2_O. *β*-estradiol (36.7 mM stock in ethanol; E2758; Sigma-Aldrich) was diluted to a final concentration of 36.7 μM in ddH_2_O. For monitoring the degradation of 1% Azo following the UV irradiation, 1% sodium 4-hexylphenylazosulfonate (Na-Azo, Excenen Pharmatech) in 100 mM TEABC buffer was 1: 200 diluted and subjected to absorbance measurement.

## Accession Numbers

Sequence data from this article can be found in the Arabidopsis Genome Initiative under the following accession numbers: *At*CNIH5 (AT4G12090), *At*PHT1;1 (AT5G43350), *At*PHT1;2 (AT5G43370), *At*PHT1;5 (AT2G32830), *At*PHT1;7 (AT3G54700), *At*PHT1;9 (AT1G76430), *At*LRK10L-1.1 (AT1G18390), *At*GXM2 (AT4G09990), *At*SOS (AT1G26650)*, At*DTX21 (AT1G33110)*, At*DTX35 (AT4G25640)*, At*URGT6 (AT1G34020), *At*OCT1 (AT1G73220), *At*HAK5 (AT4G13420), *At*VPE1 (AT3G47420), *At*SULTR1;3 (AT1G22150).

## Supporting information

**Fig. S1** Multiple marker abundance profiling (MMAP) of the identified proteins in Azo-solubilized microsomal protein extraction (MME) samples. (A) Summary of label-free proteomic analysis of microsomal proteins in *Arabidopsis* WT and *cnih5* Pi-limited roots. (B) Relative number of high confidence marker (hcm) proteins for each compartment. The relative number of distinct proteins for each compartment are displayed in the bar graph as a percentage of all unique proteins in Arabidopsis (Standard) or all proteins in the user sample (User). (C) Relative protein abundance of high hcm proteins for each compartment. The Normalized Gator Abundance Factor (NGAF) scores for each protein are summed *per* subcellular compartment according to the hcm assignments. The abundance-scaling factor for each compartment describes the skew between the standard observations in Arabidopsis (Standard) and the estimated abundance observation in the user sample. The NGAF scores for each protein in the user sample is multiplied by the compartmental scaling factor and summed per subcellular compartment according to the hcm list. The sums for each compartment are displayed in the bar graph as a percentage of the whole protein abundance (Standard) or all proteins in the total user sample (User).

**Fig. S2** Gene ontology (GO) analysis of the 4,317 identified proteins in iTRAQ-based proteomics of *Arabidopsis* WT and *cnih5* Pi-limited roots. Dot plots of the top GO terms in enrichment analysis for categories of molecular function (MF), biological process (BP), and cellular component (CC) ranked by fold enrichment. Fold enrichment compares the sample frequency representing the number of genes inputted under each GO term to the background frequency of total genes annotated to the same term in *A. thaliana*. The size of the dot represents the number of genes associated with each GO term. The more intense reddish color reflects a lower adjusted *p*-value shown by −log_10_(FDR).

**Fig. S3** GO and KEGG pathway enrichment analyses of the 372 upregulated proteins and the 51 upregulated integral microsomal proteins (MPs) in the Pi-limited root of *cnih5*. (A, B) Dot plots of the gene ontology (GO) analyses of the 372 upregulated proteins (A) and the 51 upregulated integral MPs (B) ranked by fold enrichment for categories of molecular function, biological process, and cellular component. (C, D) Dot plots of the KEGG pathway analyses of the 372 upregulated proteins (C) and the 51 upregulated integral MPs (D) ranked by fold enrichment. Fold enrichment compares the sample frequency representing the number of genes inputted under each GO term or KEGG term to the background frequency of total genes annotated to the same term in *A. thaliana*. The size of the dot indicates the number of genes in each GO term or KEGG term. The more intense reddish color reflects a lower adjusted *p*-value, as indicated by −log_10_ (FDR).

**Fig. S4** GO and KEGG pathway enrichment analyses of the 106 downregulated proteins in Pi-limited *cnih5* roots. (A, B) Dot plots of the GO (A) and KEGG pathway (B) enrichment analyses ranked by fold enrichment. Fold enrichment compares the sample frequency representing the number of genes inputted under each GO term or KEGG term to the background frequency of total genes annotated to the same term in *A. thaliana*. The size of the dot indicates the number of genes in each GO term or KEGG term. The more intense reddish color reflects a lower adj. *p*-value adjusted *p*-value shown by −log_10_(FDR).

**Fig. S5** Analysis of the interaction of PLVCub-*At*CNIH5 with NubG*-At*SOS, NubG*-At*OCT1, NubG*-At*URGT6, NubG*-At*DTX21, and NubG*-At*DTX35 using the yeast split-ubiquitin system (SUS). The co-expression of PLVCub-*At*CNIH5 with NubI and NubG are used as positive and negative controls, respectively. Yeast transformants were grown on synthetic medium without leucine and tryptophan (–LW; the left panel) or on synthetic medium lacking leucine, tryptophan, and adenine with 500 μM methionine (–LWAH; the middle panel) or on SC–LW containing 100 mg/L X-Gal (X-Gal; the right panel).

**Fig. S6** Amino acid sequence alignment of *Sc*Erv14 and *At*CNIHs. The sequence alignment was performed using the Clustal Omega algorithm (Sievers et al., 2011) and visualized using Jalview version 2.11.4.0 (Waterhouse et al., 2009). The degree of amino acid conservation among the CNIHs is represented by a light-to-dark blue color gradient. The transmembrane domains (TMD) are boxed by solid lines. The putative cornichon, YLC, IFNLL, and acidic motifs are boxed by dotted lines. The protein sequences were retrieved from UniprotKB (Uniprot ID: *Sc*Erv14, P53173; *At*CNIH1, Q9C7D7; *At*CNIH2, Q3EDD7; *At*CNIH3, Q8GWT5; *At*CNIH4, Q84W04; *At*CNIH5, Q9SZ74).

**Fig. S7** Expression of *35S:* RFP-*At*CNIH5 in *Arabidopsis*. Expression of *35S*: RFP-*At*CNIH5 in 5-day-old *Arabidopsis* seedlings grown under –P (0 µM KH_2_PO_4_, five days of starvation) conditions. Scale bars, 50 µm.

**Fig. S8** Protein expression of the endogenous *At*CNIH5 in WT and GFP-*AtCNIH5* in the *AtCNIH5pro*: GFP-*AtCNIH5/cnih5* complementation (COM) plants under Pi deficiency. Immunoblot analysis of *At*CNIH5 and GFP-*AtCNIH5* in the root of *Arabidopsis* 11-day-old WT and COM seedlings under –P (0 µM KH_2_PO_4_, seven days of starvation) conditions. The bands of *At*CNIH5 and GFP-*AtCNIH5* are indicated by solid arrowhead and open arrowhead, respectively.

**Table S1.** A list of the identified 4,317 proteins in *Arabidopsis* Pi-limited WT and *cnih5* roots with GO enrichment analysis. Detailed descriptions of our proteomic analysis, including the protein scores and mass, annotation, predicted transmembrane helices, quantification for the identified proteins, and a list of the 57 overlapped identified proteins between our datasets and the Pi starvation root proteome by Lan *et al*. (Lan et al., 2012).

**Table S2.** A list of the 372 upregulated and 106 downregulated proteins in Pi-limited *cnih5* roots with GO enrichment analysis. Detailed descriptions, including annotation, predicted transmembrane helices, and quantification for the proteins identified in proteomic analysis.

**Table S3.** Downregulated proteins in Pi-limited *cnih5* roots compared to Pi-limited WT roots by label-free proteomics across three biological replicates for each genotype.

**Table S4.** A list of the 19 root Pi starvation-induced (PSI) genes encoding transporters.

**Table S5.** Enrichment of root Pi starvation-induced (PSI) genes in the iTRAQ-based proteomic analysis of Pi-limited *cnih5* roots.

**Table S6.** Oligonucleotides used for plasmid constructs.

## Acknowledgments

This work was supported by the Ministry of Science and Technology (MOST 108-2311-B-007-003-MY3) and the National Science and Technology Council (NSTC 112-2313-B-007-001-MY3). We acknowledge Ms. He-Ting Yang for cloning the CDS of *At*OCT1, *At*HAK5, *At*VPE1, and *At*SULTR1;3, and Mr. Po-Ruey Huang for cloning the constructs expressing the truncated *At*CNIH5 variants. We also thank Mr. Cheng-Da Tsai for testing the interaction of PLVCub*-At*CNIH5 with NubG-*At*PHT1;2, NubG-*At*PHT1;5, NubG-*At*PHT1;7, and NubG-*At*PHT1;9. We thank Dr. Tzyy-Jen Chiou at Academia Sinica, Taiwan (R.O.C.), for sharing anti-*At*PHO1 and anti-*At*PHF1 antibodies and the technical support from the confocal imaging core (sponsored by MOST 110-2731-M-007-001), National Tsing Hua University.

## Author Contributions

T.-Y. L. conceived the project, designed the experiments, analyzed the experiments, and wrote the article. M.-H. T., J.-Y. W., H.-F. L., C.-Y. C., H.-X. C. and C.-A. L. performed and analyzed the experiments. T.-Y. L., M.-H. T. and J.-Y. W. contributed equally.

## References

Abas L, Luschnig C (2010) Maximum yields of microsomal-type membranes from small amounts of plant material without requiring ultracentrifugation. Analytical Biochemistry 401: 217–227

Arpat AB, Magliano P, Wege S, Rouached H, Stefanovic A, Poirier Y (2012) Functional expression of PHO1 to the Golgi and *trans*-Golgi network and its role in export of inorganic phosphate. The Plant Journal 71: 479–491

Barlowe CK, Miller EA (2013) Secretory Protein Biogenesis and Traffic in the Early Secretory Pathway. Genetics 193: 383–410

Bayle V, Arrighi J-F, Creff A, Nespoulous C, Vialaret J, Rossignol M, Gonzalez E, Paz-Ares J, Nussaume L (2011) *Arabidopsis thaliana* High-Affinity Phosphate Transporters Exhibit Multiple Levels of Posttranslational Regulation. The Plant Cell 23: 1523–1535

Bökel C, Dass S, Wilsch-Bräuninger M, Roth S (2006) *Drosophila* Cornichon acts as cargo receptor for ER export of the TGFα-like growth factor Gurken. Development 133: 459–470

Bowtell D, Fu P, Simon M, Senior P (1992) Identification of murine homologues of the Drosophila son of sevenless gene: potential activators of ras. Proceedings of the National Academy of Sciences 89: 6511–6515

Brown KA, Chen B, Guardado-Alvarez TM, Lin Z, Hwang L, Ayaz-Guner S, Jin S, Ge Y (2019) A photocleavable surfactant for top-down proteomics. Nature Methods 16: 417–420

Brown KA, Tucholski T, Eken C, Knott S, Zhu Y, Jin S, Ge Y (2020) High-Throughput Proteomics Enabled by a Photocleavable Surfactant. Angewandte Chemie International Edition 59: 8406–8410

Chen J, Wang Y, Wang F, Yang J, Gao M, Li C, Liu Y, Liu Y, Yamaji N, Ma JF, Paz-Ares J, Nussaume L, Zhang S, Yi K, Wu Z, Wu P (2015) The Rice CK2 Kinase Regulates Trafficking of Phosphate Transporters in Response to Phosphate Levels. The Plant Cell 27: 711–723

Chiu C-Y, Tsai C-D, Wang J-Y, Tsai M-H, Kanno S, Lung H-F, Liu T-Y (2024) Phosphate Starvation-Induced CORNICHON HOMOLOG 5 as Endoplasmic Reticulum Cargo Receptor for PHT1 Transporters in *Arabidopsis*. bioRxiv: 2024.2006.2020.599911

Chopin F, Wirth J, Dorbe M-F, Lejay L, Krapp A, Gojon A, Daniel-Vedele F (2007) The Arabidopsis nitrate transporter AtNRT2.1 is targeted to the root plasma membrane. Plant Physiology and Biochemistry 45: 630–635

Dancourt J, Barlowe C (2010) Protein Sorting Receptors in the Early Secretory Pathway. Annual Review of Biochemistry 79: 777–802

de Torres-Zabala M, Truman W, Bennett MH, Lafforgue G, Mansfield JW, Rodriguez Egea P, Bögre L, Grant M (2007) *Pseudomonas syringae* pv. *tomato* hijacks the *Arabidopsis* abscisic acid signalling pathway to cause disease. The EMBO Journal 26: 1434–1443

Deutsch EW, Bandeira N, Perez-Riverol Y, Sharma V, Carver Jeremy J, Mendoza L, Kundu DJ, Wang S, Bandla C, Kamatchinathan S, Hewapathirana S, Pullman Benjamin S, Wertz J, Sun Z, Kawano S, Okuda S, Watanabe Y, MacLean B, MacCoss Michael J, Zhu Y, Ishihama Y, Vizcaíno Juan A (2022) The ProteomeXchange consortium at 10 years: 2023 update. Nucleic Acids Research 51: D1539–D1548

Evenson KJ, Post-Beittenmiller D (1995) Fatty Acid-Elongating Activity in Rapidly Expanding Leek Epidermis. Plant Physiology 109: 707–716

Ge SX, Jung D, Yao R (2019) ShinyGO: a graphical gene-set enrichment tool for animals and plants. Bioinformatics 36: 2628–2629

González E, Solano R, Rubio V, Leyva A, Paz-Ares J (2005) PHOSPHATE TRANSPORTER TRAFFIC FACILITATOR1 Is a Plant-Specific SEC12-Related Protein That Enables the Endoplasmic Reticulum Exit of a High-Affinity Phosphate Transporter in *Arabidopsis*. The Plant Cell 17: 3500–3512

Hallgren J, Tsirigos KD, Pedersen MD, Almagro Armenteros JJ, Marcatili P, Nielsen H, Krogh A, Winther O (2022) DeepTMHMM predicts alpha and beta transmembrane proteins using deep neural networks. bioRxiv: 2022.2004.2008.487609

Harmel N, Cokic B, Zolles G, Berkefeld H, Mauric V, Fakler B, Stein V, Klöcker N (2012) AMPA Receptors Commandeer an Ancient Cargo Exporter for Use as an Auxiliary Subunit for Signaling. PLOS ONE 7: e30681

Hiroguchi A, Sakamoto S, Mitsuda N, Miwa K (2021) Golgi-localized membrane protein AtTMN1/EMP12 functions in the deposition of rhamnogalacturonan II and I for cell growth in Arabidopsis. Journal of Experimental Botany 72: 3611–3629

Hooper CM, Castleden IR, Tanz SK, Aryamanesh N, Millar AH (2016) SUBA4: the interactive data analysis centre for Arabidopsis subcellular protein locations. Nucleic Acids Research 45: D1064–D1074

Huang T-K, Han C-L, Lin S-I, Chen Y-J, Tsai Y-C, Chen Y-R, Chen J-W, Lin W-Y, Chen P-M, Liu T-Y, Chen Y-S, Sun C-M, Chiou T-J (2013) Identification of Downstream Components of Ubiquitin-Conjugating Enzyme PHOSPHATE2 by Quantitative Membrane Proteomics in *Arabidopsis* Roots. The Plant Cell 25: 4044–4060

Krogh A, Larsson B, von Heijne G, Sonnhammer ELL (2001) Predicting transmembrane protein topology with a hidden markov model: application to complete genomes. Journal of Molecular Biology 305: 567–580

Kurokawa K, Nakano A (2018) The ER exit sites are specialized ER zones for the transport of cargo proteins from the ER to the Golgi apparatus. The Journal of Biochemistry 165: 109–114

Lagunas-Gomez D, Yañez-Dominguez C, Zavala-Padilla G, Barlowe C, Pantoja O (2023) The C-terminus of the cargo receptor Erv14 affects COPII vesicle formation and cargo delivery. Journal of Cell Science 136

Lan P, Li W, Schmidt W (2012) Complementary Proteome and Transcriptome Profiling in Phosphate-deficient Arabidopsis Roots Reveals Multiple Levels of Gene Regulation. Molecular & Cellular Proteomics 11: 1156–1166

Lee C, Teng Q, Zhong R, Yuan Y, Haghighat M, Ye Z-H (2012) Three Arabidopsis DUF579 Domain-Containing GXM Proteins are Methyltransferases Catalyzing 4-*O*-Methylation of Glucuronic Acid on Xylan. Plant and Cell Physiology 53: 1934–1949

Lee S-B, Jung S-J, Go Y-S, Kim H-U, Kim J-K, Cho H-J, Park OK, Suh M-C (2009) Two Arabidopsis 3-ketoacyl CoA synthase genes, *KCS20* and *KCS2/DAISY*, are functionally redundant in cuticular wax and root suberin biosynthesis, but differentially controlled by osmotic stress. The Plant Journal 60: 462–475

Lelandais-Brière C, Jovanovic M, Torres GAM, Perrin Y, Lemoine R, Corre-Menguy F, Hartmann C (2007) Disruption of *AtOCT1*, an organic cation transporter gene, affects root development and carnitine-related responses in Arabidopsis. The Plant Journal 51: 154–164

Liao Y-Y, Buckhout TJ, Schmidt W (2011) Phosphate deficiency-induced cell wall remodeling. Plant Signaling & Behavior 6: 700–702

Lim CW, Yang SH, Shin KH, Lee SC, Kim SH (2015) The AtLRK10L1.2, *Arabidopsis* ortholog of wheat LRK10, is involved in ABA-mediated signaling and drought resistance. Plant Cell Reports 34: 447–455

Lin W-D, Liao Y-Y, Yang TJW, Pan C-Y, Buckhout TJ, Schmidt W (2011) Coexpression-Based Clustering of Arabidopsis Root Genes Predicts Functional Modules in Early Phosphate Deficiency Signaling. Plant Physiology 155: 1383–1402

Lin W-Y, Huang T-K, Chiou T-J (2013) NITROGEN LIMITATION ADAPTATION, a Target of MicroRNA827, Mediates Degradation of Plasma Membrane–Localized Phosphate Transporters to Maintain Phosphate Homeostasis in *Arabidopsis*. The Plant Cell 25: 4061–4074

Liu T-Y (2021) Using Tripartite Split-sfGFP for the Study of Membrane Protein–Protein Interactions. *In* JJ Sanchez-Serrano, J Salinas, eds, Arabidopsis Protocols. Springer US, New York, NY, pp 323–336

Liu T-Y, Chou W-C, Chen W-Y, Chu C-Y, Dai C-Y, Wu P-Y (2018) Detection of membrane protein–protein interaction *in planta* based on dual-intein-coupled tripartite split-GFP association. The Plant Journal 94: 426–438

Liu T-Y, Huang T-K, Tseng C-Y, Lai Y-S, Lin S-I, Lin W-Y, Chen J-W, Chiou T-J (2012) PHO2-Dependent Degradation of PHO1 Modulates Phosphate Homeostasis in *Arabidopsis*. The Plant Cell 24: 2168–2183

Liu T-Y, Huang T-K, Yang S-Y, Hong Y-T, Huang S-M, Wang F-N, Chiang S-F, Tsai S-Y, Lu W-C, Chiou T-J (2016) Identification of plant vacuolar transporters mediating phosphate storage. Nature Communications 7: 11095

Mølhøj M, Verma R, Reiter W-D (2004) The Biosynthesis of d-Galacturonate in Plants. Functional Cloning and Characterization of a Membrane-Anchored UDP-d-Glucuronate 4-Epimerase from Arabidopsis. Plant Physiology 135: 1221–1230

Mouille G, Ralet M-C, Cavelier C, Eland C, Effroy D, Hématy K, McCartney L, Truong HN, Gaudon V, Thibault J-F, Marchant A, Höfte H (2007) Homogalacturonan synthesis in *Arabidopsis thaliana* requires a Golgi-localized protein with a putative methyltransferase domain. The Plant Journal 50: 605–614

Nakanishi H, Suda Y, Neiman AM (2007) Erv14 family cargo receptors are necessary for ER exit during sporulation in *Saccharomyces cerevisiae*. Journal of Cell Science 120: 908–916

Nguyen TV, Gupta R, Annas D, Yoon J, Kim Y-J, Lee GH, Jang JW, Park KH, Rakwal R, Jung K-H, Min CW, Kim ST (2021) An Integrated Approach for the Efficient Extraction and Solubilization of Rice Microsomal Membrane Proteins for High-Throughput Proteomics. Frontiers in Plant Science 12

Nussaume L, Kanno S, Javot H, Marin E, Nakanishi TM, Thibaud M-C (2011) Phosphate Import in Plants: Focus on the PHT1 Transporters. Frontiers in Plant Science 2

Obrdlik P, El-Bakkoury M, Hamacher T, Cappellaro C, Vilarino C, Fleischer C, Ellerbrok H, Kamuzinzi R, Ledent V, Blaudez D, Sanders D, Revuelta JL, Boles E, André B, Frommer WB (2004) K^+^ channel interactions detected by a genetic system optimized for systematic studies of membrane protein interactions. Proceedings of the National Academy of Sciences 101: 12242–12247

Pagant S, Wu A, Edwards S, Diehl F, Miller Elizabeth A (2015) Sec24 Is a Coincidence Detector that Simultaneously Binds Two Signals to Drive ER Export. Current Biology 25: 403–412

Park BS, Seo JS, Chua N-H (2014) NITROGEN LIMITATION ADAPTATION Recruits PHOSPHATE2 to Target the Phosphate Transporter PT2 for Degradation during the Regulation of *Arabidopsis* Phosphate Homeostasis. The Plant Cell 26: 454–464

Perez-Riverol Y, Bandla C, Kundu Deepti J, Kamatchinathan S, Bai J, Hewapathirana S, John Nithu S, Prakash A, Walzer M, Wang S, Vizcaíno Juan A (2024) The PRIDE database at 20 years: 2025 update. Nucleic Acids Research 53: D543–D553

Poirier Y, Bucher M (2002) Phosphate Transport and Homeostasis in Arabidopsis. The Arabidopsis Book 2002

Poirier Y, Jaskolowski A, Clúa J (2022) Phosphate acquisition and metabolism in plants. Current Biology 32: R623–R629

Powers J, Barlowe C (1998) Transport of Axl2p Depends on Erv14p, an ER–Vesicle Protein Related to the *Drosophila cornichon* Gene Product. Journal of Cell Biology 142: 1209–1222

Powers J, Barlowe C (2002) Erv14p Directs a Transmembrane Secretory Protein into COPII-coated Transport Vesicles. Molecular Biology of the Cell 13: 880–891

R Core Team (2021) R: A Language and Environment for Statistical Computing. R Foundation for Statistical Computing

Ramel F, Sulmon C, Cabello-Hurtado F, Taconnat L, Martin-Magniette M-L, Renou J-P, El Amrani A, Couée I, Gouesbet G (2007) Genome-wide interacting effects of sucrose and herbicide-mediated stress in *Arabidopsis thaliana*: novel insights into atrazine toxicity and sucrose-induced tolerance. BMC Genomics 8: 450

Rosas-Santiago P, Lagunas-Gómez D, Barkla BJ, Vera-Estrella R, Lalonde S, Jones A, Frommer WB, Zimmermannova O, Sychrová H, Pantoja O (2015) Identification of rice cornichon as a possible cargo receptor for the Golgi-localized sodium transporter OsHKT1;3. Journal of Experimental Botany 66: 2733–2748

Rosas-Santiago P, Lagunas-Gomez D, Yáñez-Domínguez C, Vera-Estrella R, Zimmermannová O, Sychrová H, Pantoja O (2017) Plant and yeast cornichon possess a conserved acidic motif required for correct targeting of plasma membrane cargos. Biochimica et Biophysica Acta (BBA) - Molecular Cell Research 1864: 1809–1818

Roth S, Shira Neuman-Silberberg F, Barcelo G, Schüpbach T (1995) cornichon and the EGF receptor signaling process are necessary for both anterior-posterior and dorsal-ventral pattern formation in Drosophila. Cell 81: 967–978

RStudio Team (2020) RStudio: Integrated Development Environment for R. RStudio, PBC

Saez-Aguayo S, Parra-Rojas JP, Sepúlveda-Orellana P, Celiz-Balboa J, Arenas-Morales V, Sallé C, Salinas-Grenet H, Largo-Gosens A, North HM, Ralet M-C, Orellana A (2020) Transport of UDP-rhamnose by URGT2, URGT4, and URGT6 modulates rhamnogalacturonan-I length. Plant Physiology 185: 914–933

Salazar-Henao JE, Schmidt W (2016) An Inventory of Nutrient-Responsive Genes in *Arabidopsis* Root Hairs. Frontiers in Plant Science 7

Sievers F, Wilm A, Dineen D, Gibson TJ, Karplus K, Li W, Lopez R, McWilliam H, Remmert M, Söding J, Thompson JD, Higgins DG (2011) Fast, scalable generation of high-quality protein multiple sequence alignments using Clustal Omega. Molecular Systems Biology 7: 539

Slowikowski K, Schep A, Hughes S, Kien Dang T, Lukauskas S, Irisson J-O, Kamvar ZN, Ryan T, Christophe D, Hiroaki Y, Gramme P, Abdol AM, Barrett M, Cannoodt R, Krassowski M, Chirico M, Aphalo P, Barton F (2024) ggrepel: Automatically Position Non-Overlapping Text Labels with ’ggplot2’. In,

Spitzer M, Wildenhain J, Rappsilber J, Tyers M (2014) BoxPlotR: a web tool for generation of box plots. Nature Methods 11: 121–122

Sterling JD, Atmodjo MA, Inwood SE, Kumar Kolli VS, Quigley HF, Hahn MG, Mohnen D (2006) Functional identification of an *Arabidopsis* pectin biosynthetic homogalacturonan galacturonosyltransferase. Proceedings of the National Academy of Sciences 103: 5236–5241

The UniProt Consortium (2023) UniProt: the Universal Protein Knowledgebase in 2023. Nucleic Acids Research 51: D523–D531

Tian T, Liu Y, Yan H, You Q, Yi X, Du Z, Xu W, Su Z (2017) agriGO v2.0: a GO analysis toolkit for the agricultural community, 2017 update. Nucleic Acids Research 45: W122–W129

Urbanowicz BR, Peña MJ, Ratnaparkhe S, Avci U, Backe J, Steet HF, Foston M, Li H, O’Neill MA, Ragauskas AJ, Darvill AG, Wyman C, Gilbert HJ, York WS (2012) 4-*O*-methylation of glucuronic acid in *Arabidopsis* glucuronoxylan is catalyzed by a domain of unknown function family 579 protein. Proceedings of the National Academy of Sciences 109: 14253–14258

Wang X, Liang S (2012) Sample preparation for the analysis of membrane proteomes by mass spectrometry. Protein & Cell 3: 661–668

Waterhouse AM, Procter JB, Martin DMA, Clamp M, Barton GJ (2009) Jalview Version 2—a multiple sequence alignment editor and analysis workbench. Bioinformatics 25: 1189–1191

Wei L, Liu L, Chen Z, Huang Y, Yang L, Wang P, Xue S, Bie Z (2023) CmCNIH1 improves salt tolerance by influencing the trafficking of CmHKT1;1 in pumpkin. The Plant Journal 114: 1353–1368

Wickham H (2023) stringr: Simple, Consistent Wrappers for Common String Operations. In,

Wickham H, Chang W, Henry L, Lin Pedersen T, Takahashi K, Wilke C, Woo K, Yutani H, Dunnington D, van den Brand T (2016) ggplot2: Elegant Graphics for Data Analysis. In. Springer-Verlag New York

Wu X, Lai Y, Rao S, Lv L, Ji M, Han K, Weng J, Lu Y, Peng J, Lin L, Wu G, Chen J, Yan F, Zheng H (2021) Genome-Wide Identification Reveals That *Nicotiana benthamiana* Hypersensitive Response (HR)-Like Lesion Inducing Protein 4 (NbHRLI4) Mediates Cell Death and Salicylic Acid-Dependent Defense Responses to Turnip Mosaic Virus. Frontiers in Plant Science 12

Wudick MM, Portes MT, Michard E, Rosas-Santiago P, Lizzio MA, Nunes CO, Campos C, Santa Cruz Damineli D, Carvalho JC, Lima PT, Pantoja O, Feijó JA (2018) CORNICHON sorting and regulation of GLR channels underlie pollen tube Ca^2+^ homeostasis. Science 360: 533–536

Yáñez-Domínguez C, Lagunas-Gómez D, Torres-Cifuentes DM, Bezanilla M, Pantoja O (2023) A cornichon protein controls polar localization of the PINA auxin transporter in *Physcomitrium patens*. Development 150

Yang Z, Yang J, Wang Y, Wang F, Mao W, He Q, Xu J, Wu Z, Mao C (2020) PROTEIN PHOSPHATASE95 Regulates Phosphate Homeostasis by Affecting Phosphate Transporter Trafficking in Rice. The Plant Cell 32: 740–757

Yoshimoto K, Hanaoka H, Sato S, Kato T, Tabata S, Noda T, Ohsumi Y (2004) Processing of ATG8s, Ubiquitin-Like Proteins, and Their Deconjugation by ATG4s Are Essential for Plant Autophagy. The Plant Cell 16: 2967–2983

Yuan Y, Teng Q, Lee C, Zhong R, Ye Z-H (2014) Modification of the degree of 4-*O*-methylation of secondary wall glucuronoxylan. Plant Science 219–220: 42-50

Zhang H, Guo Z, Zhuang Y, Suo Y, Du J, Gao Z, Pan J, Li L, Wang T, Xiao L, Qin G, Jiao Y, Cai H, Li L (2021) MicroRNA775 regulates intrinsic leaf size and reduces cell wall pectin levels by targeting a galactosyltransferase gene in Arabidopsis. The Plant Cell 33: 581–602

Zhang H, Zhao F-G, Tang R-J, Yu Y, Song J, Wang Y, Li L, Luan S (2017) Two tonoplast MATE proteins function as turgor-regulating chloride channels in *Arabidopsis*. Proceedings of the National Academy of Sciences 114: E2036–E2045

Zheng J, Yao L, Zeng X, Wang B, Pan L (2023) *ERV14* receptor impacts mycelial growth via its interactions with cell wall synthase and transporters in *Aspergillus niger*. Frontiers in Microbiology 14

Zhou M, Zhu S, Mo X, Guo Q, Li Y, Tian J, Liang C (2022) Proteomic Analysis Dissects Molecular Mechanisms Underlying Plant Responses to Phosphorus Deficiency. Cells 11: 651

Zimmermannová O, Felcmanová K, Rosas-Santiago P, Papoušková K, Pantoja O, Sychrová H (2019) Erv14 cargo receptor participates in regulation of plasma-membrane potential, intracellular pH and potassium homeostasis via its interaction with K^+^-specific transporters Trk1 and Tok1. Biochimica et Biophysica Acta (BBA) - Molecular Cell Research 1866: 1376–1388

