## Supplemental data for "CORNICHON HOMOLOG 5-dependent ER export of membrane cargoes in phosphate-starved *Arabidopsis* root as revealed by membrane proteomic analysis"

A

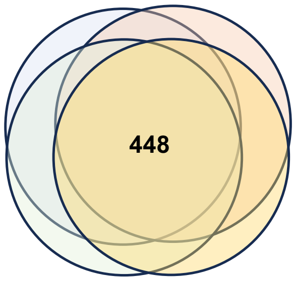


| Sample | No. of identified proteins |
|---|---|
| WT-1 | 593 |
| WT-2 | 721 |
| *cnih5*-1 | 660 |
| *cnih5*-2 | 686 |

B

C

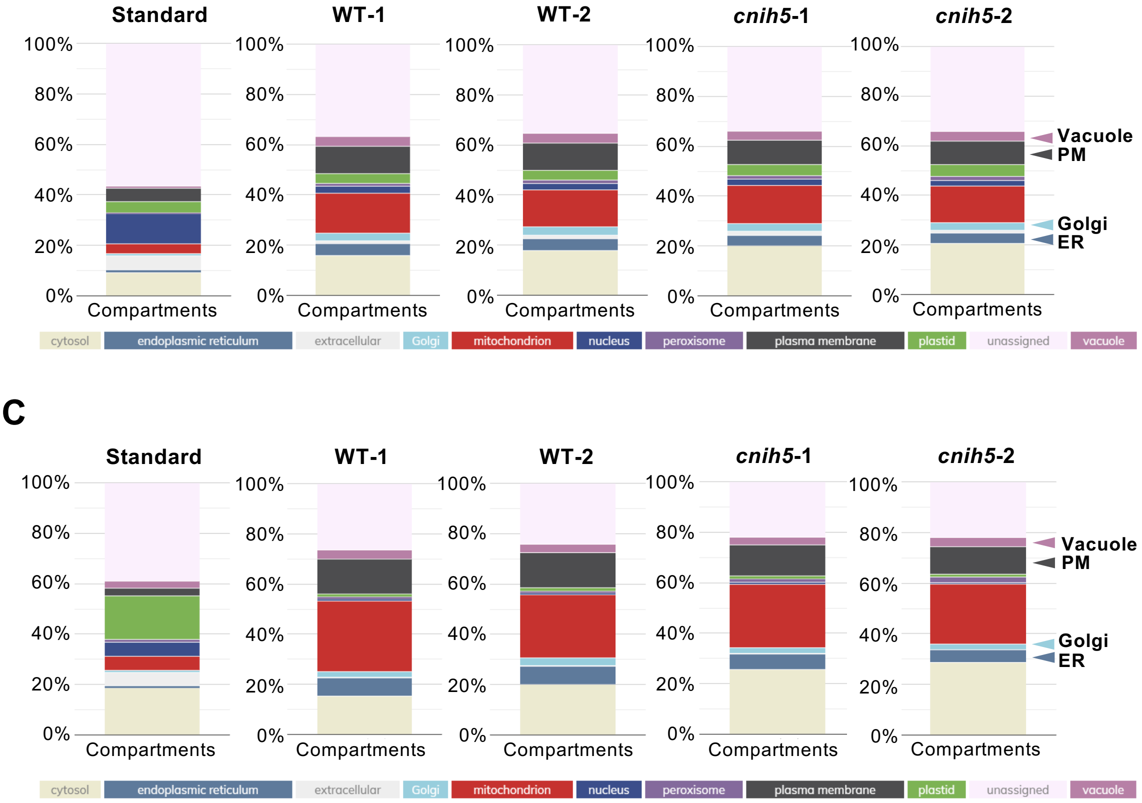


**Fig. S1** Multiple marker abundance profiling (MMAP) of the identified proteins in Azo-solubilized microsomal protein extraction (MME) samples. (A) Summary of label-free proteomic analysis of microsomal proteins in *Arabidopsis* WT and *cnih5* Pi-limited roots. (B) Relative number of high confidence marker (hcm) proteins for each compartment. The relative number of distinct proteins for each compartment are displayed in the bar graph as a percentage of all unique proteins in Arabidopsis (Standard) or all proteins in the user sample (User). (C) Relative protein abundance of high hcm proteins for each compartment. The Normalized Gator Abundance Factor (NGAF) scores for each protein are summed *per* subcellular compartment according to the hcm assignments. The abundance-scaling factor for each compartment describes the skew between the standard observations in Arabidopsis (Standard) and the estimated abundance observation in the user sample. The NGAF scores for each protein in the user sample is multiplied by the compartmental scaling factor and summed per subcellular compartment according to the hcm list. The sums for each compartment are displayed in the bar graph as a percentage of the whole protein abundance (Standard) or all proteins in the total user sample (User).

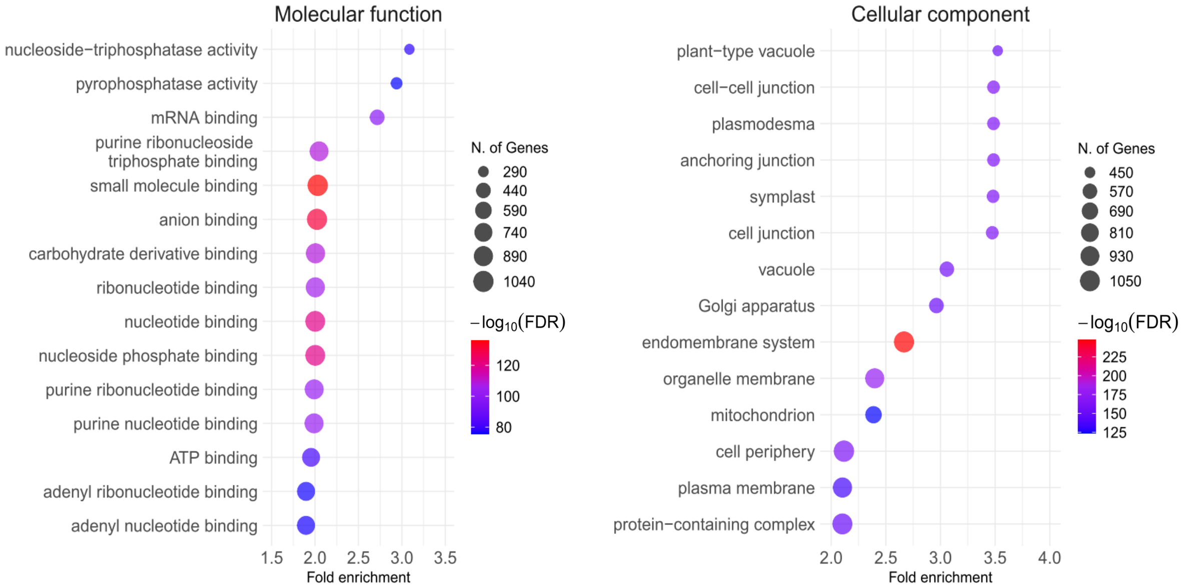


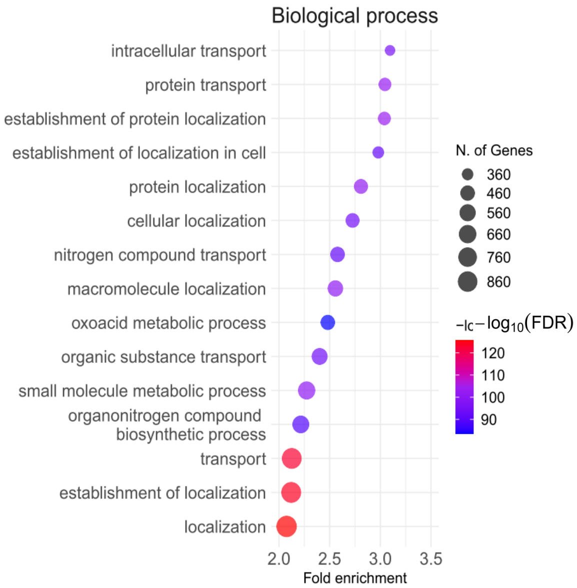


**Fig. S2** Gene ontology (GO) analysis of the 4,317 identified proteins in iTRAQ-based proteomics of *Arabidopsis* WT and *cnih5* Pi-limited roots. Dot plots of the top GO terms in enrichment analysis for categories of molecular function (MF), biological process (BP), and cellular component (CC) ranked by fold enrichment. Fold enrichment compares the sample frequency representing the number of genes inputted under each GO term to the background frequency of total genes annotated to the same term in *A. thaliana*. The size of the dot represents the number of genes associated with each GO term. The more intense reddish color reflects a lower adjusted *p*-value shown by $-\log_{10}$(FDR).

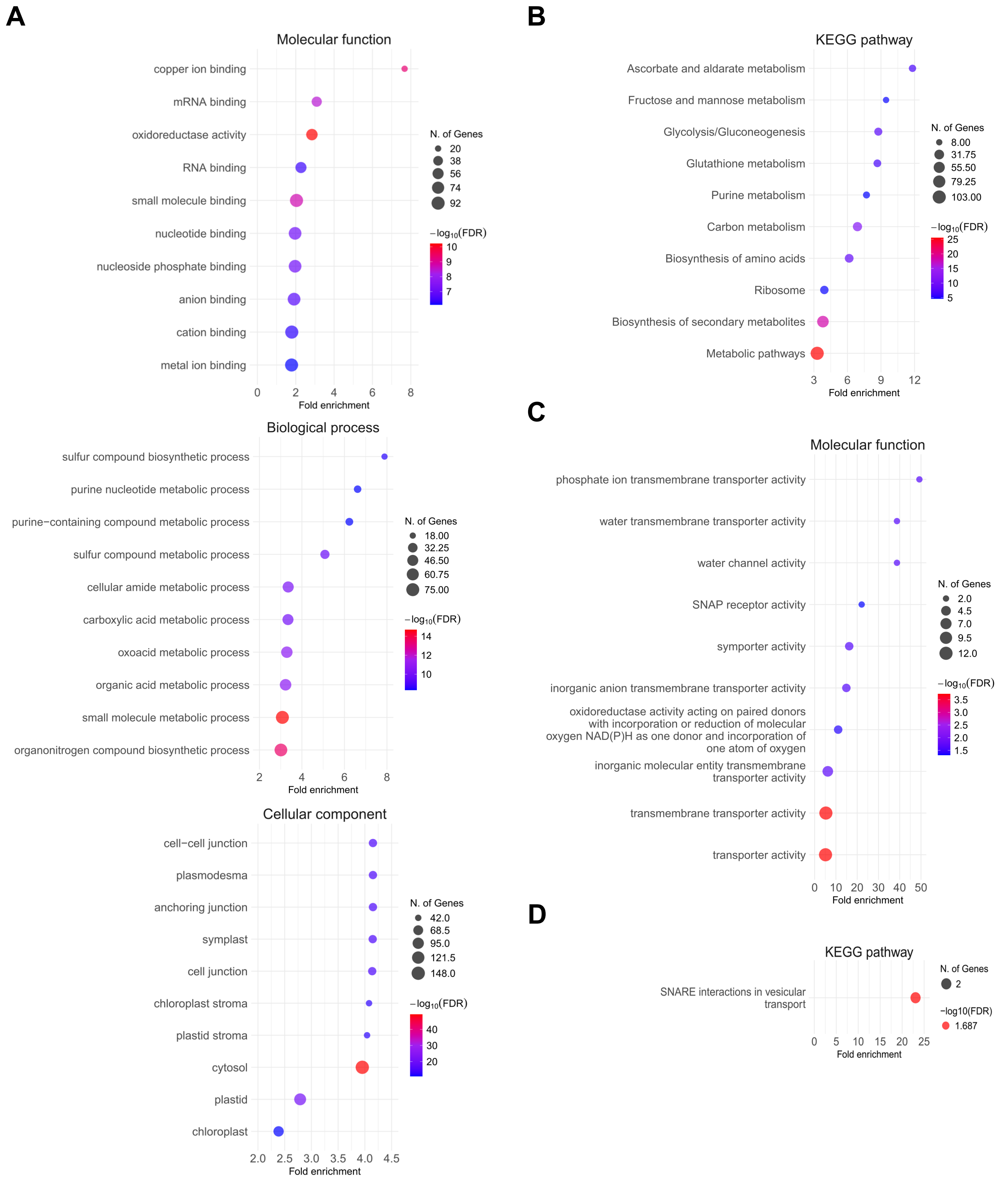


**Fig. S3** GO and KEGG pathway enrichment analyses of the 372 upregulated proteins and the 51 upregulated integral microsomal proteins (MPs) in the Pi-limited root of *cnih5*. (A, B) Dot plots of the gene ontology (GO) analyses of the 372 upregulated proteins (A) and the 51 upregulated integral MPs (B) ranked by fold enrichment for categories of molecular function, biological process, and cellular component. (C, D) Dot plots of the KEGG pathway analyses of the 372 upregulated proteins (C) and the 51 upregulated integral MPs (D) ranked by fold enrichment. Fold enrichment compares the sample frequency representing the number of genes inputted under each GO term or KEGG term to the background frequency of total genes annotated to the same term in *A. thaliana*. The size of the dot indicates the number of genes in each GO term or KEGG term. The more intense reddish color reflects a lower adjusted *p*-value, as indicated by $-\log_{10}$ (FDR).

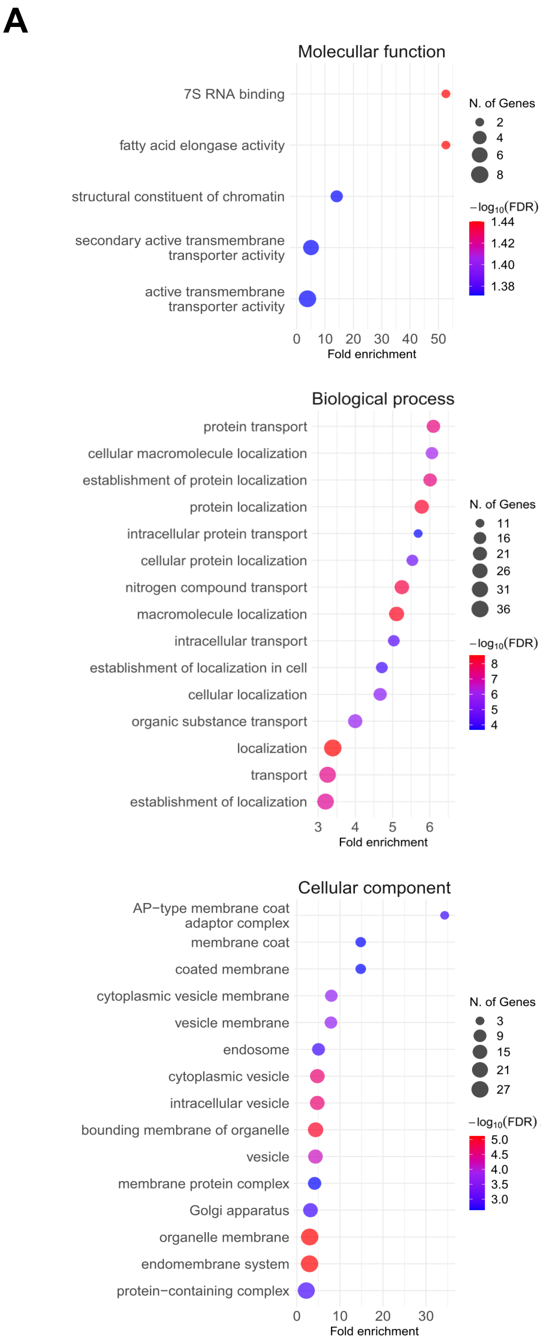


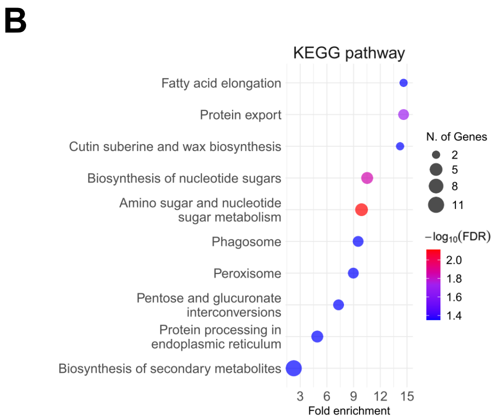


**Fig. S4** GO and KEGG pathway enrichment analyses of the 106 downregulated proteins in Pi-limited *cnih5* roots. (A, B) Dot plots of the GO (A) and KEGG pathway (B) enrichment analyses ranked by fold enrichment. Fold enrichment compares the sample frequency representing the number of genes inputted under each GO term or KEGG term to the background frequency of total genes annotated to the same term in *A. thaliana*. The size of the dot indicates the number of genes in each GO term or KEGG term. The more intense reddish color reflects a lower adj. *p*-value adjusted *p*-value shown by $-\log_{10}$(FDR).

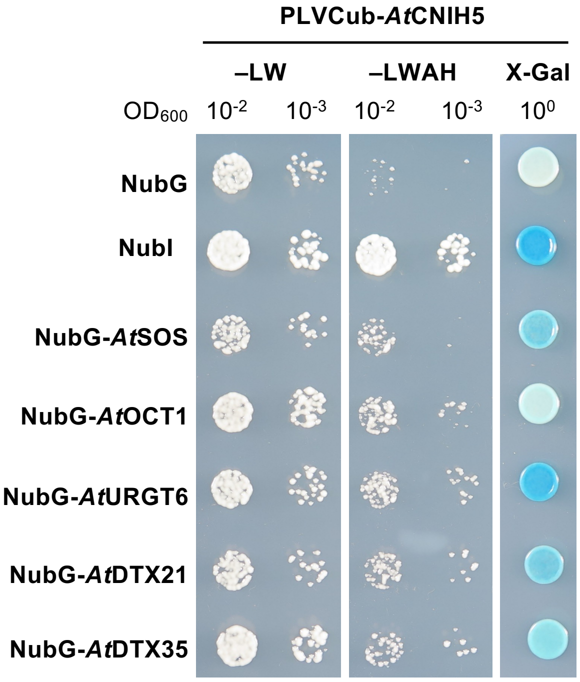


**Fig. S5** Analysis of the interaction of PLVCub-*At*CNIH5 with NubG-*At*SOS, NubG-*At*OCT1, NubG-*At*URGT6, NubG-*At*DTX21, and NubG-*At*DTX35 using the yeast split-ubiquitin system (SUS). The co-expression of PLVCub-*At*CNIH5 with NubI and NubG are used as positive and negative controls, respectively. Yeast transformants were grown on synthetic medium without leucine and tryptophan (–LW; the left panel) or on synthetic medium lacking leucine, tryptophan, and adenine with 500 μM methionine (–LWAH; the middle panel) or on SC–LW containing 100 mg/L X-Gal (X-Gal; the right panel).

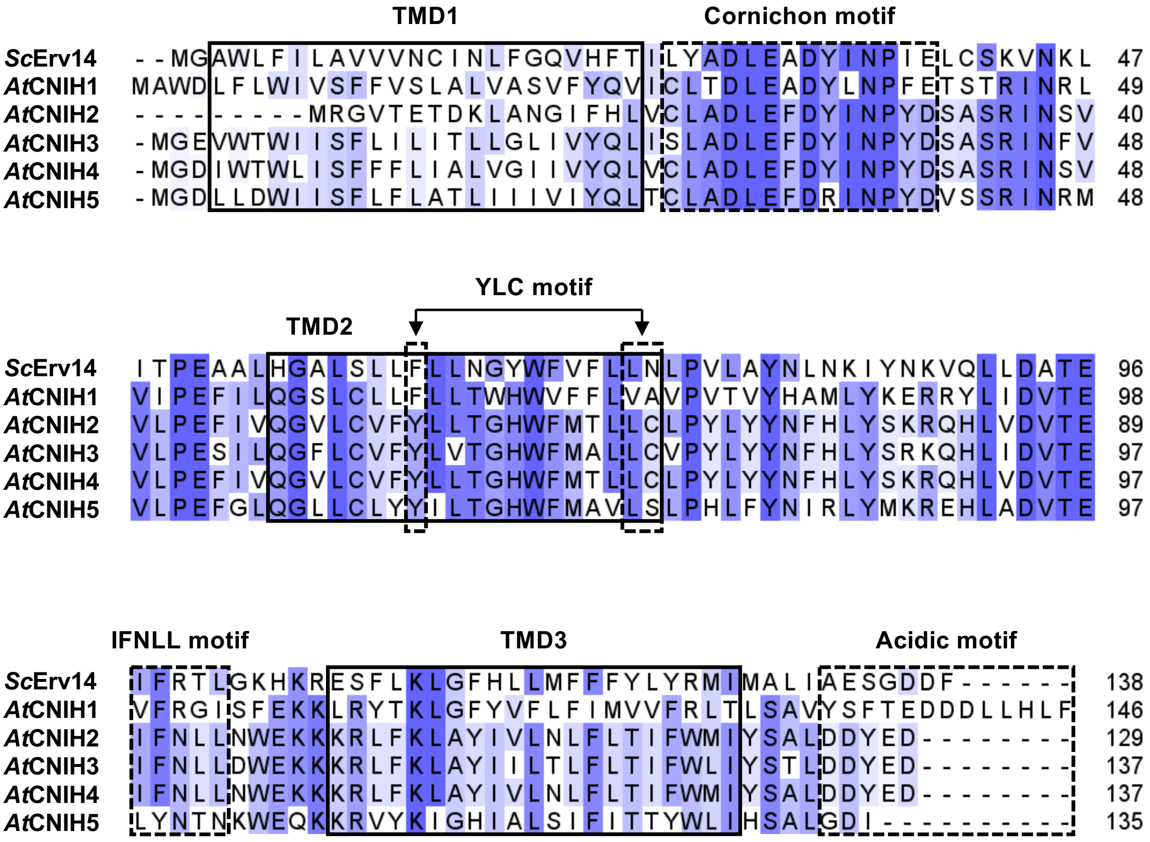


**Fig. S6** Amino acid sequence alignment of *Sc*Erv14 and *At*CNIHs. The sequence alignment was performed using the Clustal Omega algorithm and visualized using Jalview version 2.11.4.0 . The degree of amino acid conservation among the CNIHs is represented by a light-to-dark blue color gradient. The transmembrane domains (TMD) are boxed by solid lines. The putative cornichon, YLC, IFNLL, and acidic motifs are boxed by dotted lines. The protein sequences were retrieved from UniprotKB (Uniprot ID: *Sc*Erv14, P53173; *At*CNIH1, Q9C7D7; *At*CNIH2, Q3EDD7; *At*CNIH3, Q8GWT5; *At*CNIH4, Q84W04; *At*CNIH5, Q9SZ74).

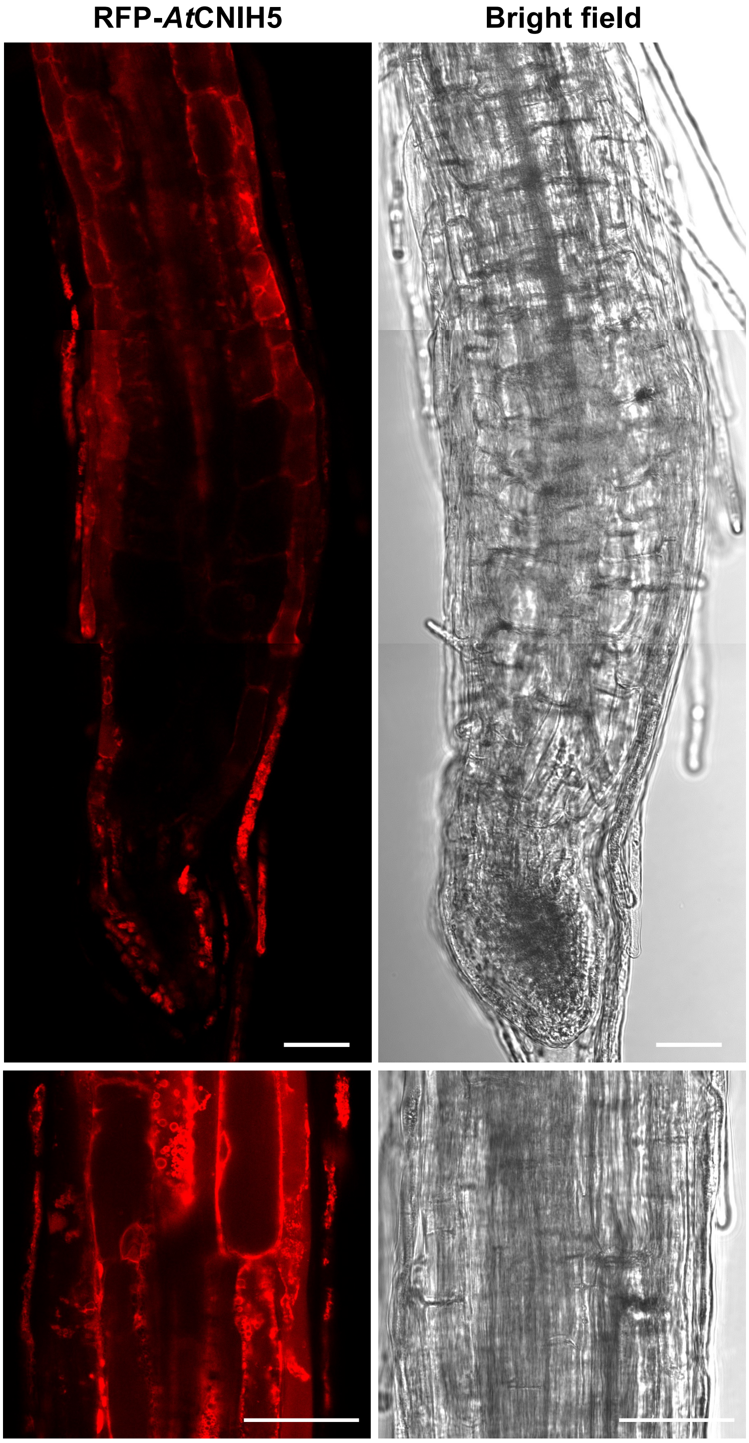


**Fig. S7** Expression of *35S:* RFP-*At*CNIH5 in *Arabidopsis*. Expression of *35S*: RFP-*At*CNIH5 in 5-day-old *Arabidopsis* seedlings grown under –P (0 μM $KH_2PO_4$, five days of starvation) conditions. Scale bars, 50 μm.

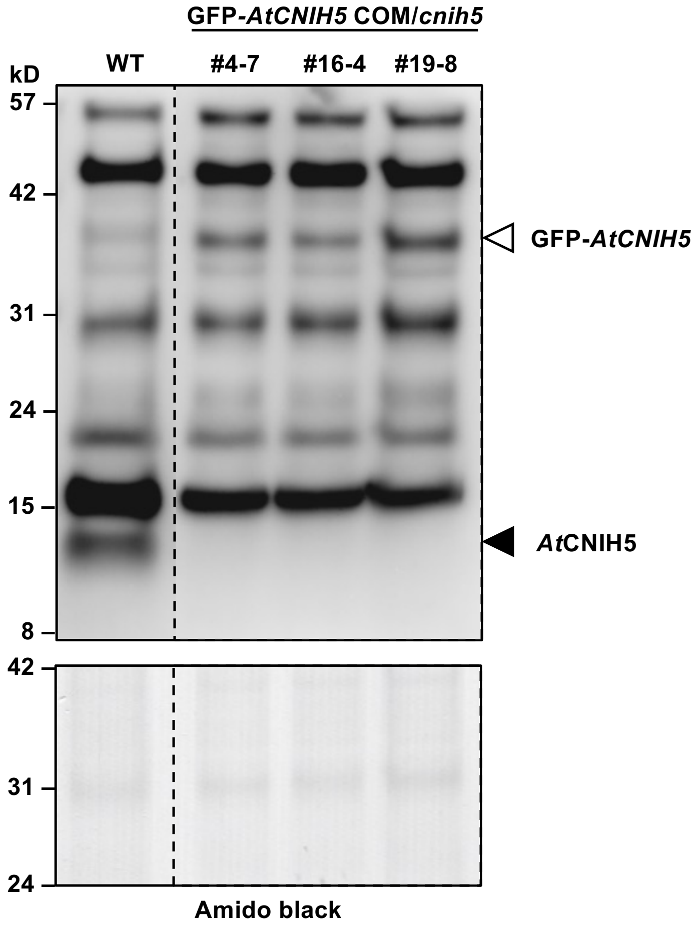


**Fig. S8** Protein expression of the endogenous *At*CNIH5 in WT and GFP-*AtCNIH5* in the *AtCNIH5pro*: GFP-*AtCNIH5/cnih5* complementation (COM) plants under Pi deficiency. Immunoblot analysis of *At*CNIH5 and GFP-*AtCNIH5* in the root of *Arabidopsis* 11-day-old WT and COM seedlings under –P (0 µM $KH_2PO_4$, seven days of starvation) conditions. The bands of *At*CNIH5 and GFP-*AtCNIH5* are indicated by solid arrowhead and open arrowhead, respectively.

**Supplementary Table S3.** Downregulated proteins in Pi-limited *cnih5* roots compared to Pi-limited WT roots by label-free proteomics across three biological replicates for each genotype. Relative protein abundance (%), defined by identified peptide spectrum counting (#PSM) divided by the total number of #PSM in a sample with a false discovery rate (FDR) of 1% x100. CV, coefficient of variation defined as the ratio of the standard deviation to the mean of relative protein abundance. Normalized ratio is defined by the protein abundance ratio divided by geometric median of protein group ratios in each group comparison. Only proteins showing a consistent trend of downregulation between the label-free (normalized ratio ≤ 0.82) and the iTRAQ-labeling (normalized ratio ≤ 0.75) methods in this study are listed. *At*OCT1, *At*DTX35, *At*TIP2;2, and *At*KCS20 showed a weak trend of downregulation (normalized ratios 0.88−0.92). n.d., not detected.

| AGI code | Accession | Gene name | Relative protein abundance (%) | | | | | | | | Normalized ratio (adj. *p*-value) |
|---|---|---|---|---|---|---|---|---|---|---|---|
| | | | WT−P _1 | WT −P _2 | WT −P _3 | CV | *cnih5* −P _1 | *cnih5* −P _2 | *cnih5* −P _3 | CV | *cnih5* −P/ WT −P |
| AT5G43350 | Q8VYM2 | Inorganic phosphate transporter 1;1 (*AtPHT1;1*) | 0.26 | 0.26 | 0.27 | 2.6 | 0.24 | 0.21 | 0.21 | 6.7 | 0.77 (1.24E-01) |
| AT5G43370 | Q96243 | Inorganic phosphate transporter 1;2 (*AtPHT1;2*) | 0.25 | 0.24 | 0.26 | 3.6 | 0.21 | 0.20 | 0.19 | 5.9 | 0.72 (1.41E-03) |
| AT5G43360 | O48639 | Inorganic phosphate transporter 1;3 (*AtPHT1;3*) | 0.06 | 0.07 | 0.07 | 8.9 | n.d. | n.d. | n.d. | - | 0.33 (8.93E-16) |
| AT2G38940 | Q96303 | Inorganic phosphate transporter 1;4 (*AtPHT1;4*) | 0.04 | 0.04 | 0.05 | 11.4 | 0.02 | 0.03 | 0.03 | 13.5 | 0.67 (4.02E-05) |
| AT4G30440 | Q9M0B6 | UDP-D-glucuronate 4-epimerase 1 (*AtGAE1*) | - | 0.02 | 0.01 | 17.7 | n.d. | 0.01 | n.d. | - | 0.71 (1.77E-01) |
| AT4G01630 | Q9ZSI1 | EXPANSIN 17 (*AtEXPA17*) | 0.01 | 0.02 | 0.01 | 14.2 | 0.01 | n.d. | n.d. | - | 0.82 (2.27E-01) |
| AT4G09990 | Q9T0F7 | Glucuronoxylan methyltransferase 2 (*AtGXM2*) | 0.02 | 0.02 | 0.02 | 15.0 | n.d. | 0.02 | 0.01 | 12.4 | 0.79 (2.95E-01) |
| AT2G17380 | Q8LEZ8 | CLATHRIN ASSEMBLY PROTEIN AP19; AP-1 complex subunit sigma-1 (*AtAAP19-1*) | 0.02 | 0.01 | 0.01 | 32.7 | 0.01 | 0.01 | n.d. | 5.1 | 0.78 (2.73E-01) |

**Supplementary Table S4.** List of 19 root Pi starvation-induced (PSI) genes encoding transporters.

| Accession No. | RPKM values | | | $Log_2FC$ of RPKM values | | | Description |
|---|---|---|---|---|---|---|---|
| | P0 | P1 | P3 | P1/P0 | P3/P0 | P3/P0−P1/P0 | |
| AT5G43360 | 0.54 | 1.04 | 53.88 | 0.96 | 6.65 | 5.69 | PHT1;3 |
| AT1G73220 | 1.07 | 8.33 | 91.17 | 2.96 | 6.41 | 3.45 | OCT1 |
| AT5G43370 | 4.24 | 16.44 | 186.53 | 1.96 | 5.46 | 3.50 | PHT1;2 |
| AT1G20860 | 0.25 | 0.39 | 10.08 | 0.63 | 5.33 | 4.70 | PHT1;8 |
| AT3G47420 | 10.61 | 47.73 | 240.43 | 2.17 | 4.50 | 2.33 | VPE1 |
| AT1G22150 | 0.25 | 0.97 | 4.95 | 1.96 | 4.31 | 2.35 | SULTR1;3 |
| AT1G76430 | 0.74 | 1.37 | 13.32 | 0.88 | 4.16 | 3.28 | PHT1;9 |
| AT4G13420 | 0.25 | 0.53 | 2.61 | 1.09 | 3.39 | 2.29 | HAK5 |
| AT2G38940 | 20.24 | 53.35 | 205.21 | 1.40 | 3.34 | 1.94 | PHT1;4 |
| AT4G15540 | 12.82 | 25.94 | 73.79 | 1.02 | 2.52 | 1.51 | EamA-like transporter |
| AT1G11460 | 4.17 | 9.12 | 17.18 | 1.13 | 2.04 | 0.91 | EamA-like transporter |
| AT3G58060 | 0.28 | 0.55 | 1.16 | 0.96 | 2.04 | 1.08 | MTP8 |
| AT5G48485 | 14.62 | 39.19 | 54.76 | 1.42 | 1.91 | 0.48 | DIR1 |
| AT5G43350 | 216.27 | 349.43 | 712.94 | 0.69 | 1.72 | 1.03 | PHT1;1 |
| AT3G51860 | 9.68 | 16.47 | 27.72 | 0.77 | 1.52 | 0.75 | CAX3 |
| AT3G01760 | 1.57 | 2.26 | 4.01 | 0.52 | 1.35 | 0.83 | LHT6 |
| AT1G22530 | 67.83 | 98.91 | 144.30 | 0.54 | 1.09 | 0.54 | PATL2 |
| AT2G22500 | 18.06 | 27.48 | 38.02 | 0.61 | 1.07 | 0.47 | DIC1 |
| AT5G19410 | 7.26 | 10.76 | 15.20 | 0.57 | 1.07 | 0.50 | ABCG23 |

RNA-seq data derived from Liu *et al.*, 2016

**Supplementary Table S5.** Enrichment of root Pi starvation-induced (PSI) genes in the iTRAQ-based proteomic analysis

| Categories | No. of proteins | [a]No. of root PSI genes | Enrichment of root PSi genes (%) |
|---|---|---|---|
| Total identified proteins | 4317 | 252 | 5.8 |
| Upregulated proteins | 372 | 32 | 8.6 |
| Upregulated integral MPs | 51 | 4 | 7.8 |
| Downregulated proteins | 106 | 15 | 14.2 |
| Downregulated integral MPs | 31 | 10 | 32.3 |

[a]PSI genes were determined by FC ≥ 1.5 (P3/P0) in RNA-seq data from Liu *et al.*, 2016.

**Supplementary Table S6.** Oligonucleotides used for plasmid constructs.

| | | |
|---|---|---|
| **Yeast expression vectors** | | |
| PLVCub-*At*CNIH5 | EcoRI_*At*CNIH5.for | gaattcATGGGCGATCTACTCGATTGGATTA |
| | *At*CNIH5 STOP_EcoRI.rev | gaattcTTAGATGTCTCCTAATGCCGAGTGG |
| *At*LRK10L1.1- NubG | EcoRI_*At*LRK10L1.1.for | aagaattcATGTCCATCTTTTTTTTCTTCATCAG |
| | *At*LRK10L1.1 (noSTOP)_EcoRI.rev | aagaattcCTTACTGTCCCATTTCACAATG |
| NubG-*At*SOS/ *At*SOS-NubG | MunI_*At*SOS.for | aacaattgATGGAGACGGAAACGAATCAGG |
| | *At*SOS (noSTOP)_MunI.rev | aacaattgCTCAGCATCCACAACCGTAAC |
| NubG-*At*DTX21/ *At*DTX21-NubG | MunI_*At*DTX21.for | ttcaattgATGGCCGGAGGAGGAGGAGA |
| | *At*DTX21 (noSTOP)_MunI.rev | tacaattgTTCCTCTGATGATACTTGGTTTACGTCTC |
| NubG-*At*URGT6/ *At*URGT6-NubG | EcoRI_*At*URGT6.for | aagaattcATGGCTCCAGTGAGTAAAGC |
| | *At*URGT6 (noSTOP)_EcoRI.rev | aagaattcGGCTTTGTCTTCGTTGTCGTC |
| NubG-*At*OCT1/ *At*OCT1-NubG | MunI_*At*OCT1.for | aacaattgATGGAACCTTCAAAACAAGAAG |
| | *At*OCT1 (noSTOP)_MunI.rev | aacaattgAGTAATCATGATTGTTTCGTTTTC |
| NubG-*At*GXM2 | EcoRI_*At*GXM2.for | aagaattcATGAGGAATAAATCCCAATCATTCATC |
| | *At*GXM2 (noSTOP)_EcoRI.rev | aagaattcAAAGCGGCGACTGATATCCG |
| NubG-*At*DTX35/ *At*DTX35-NubG | EcoRI_*At*DTX35.for | aagaattcATGGATCCGACGGCGCCGTT |
| | *At*DTX35 (noSTOP)_EcoRI.rev | aagaattcCGCAAGTATATCCTTGGATGTCGTCTCAC |
| NubG-*At*CNIH5 | EcoRI_*At*CNIH5.for | gaattcATGGGCGATCTACTCGATTGGATTA |
| | *At*CNIH5 noSTOP_EcoRI.rev | gaattcGATGTCTCCTAATGCCGAGTGGATC |
| NubG-*At*CNIH5 1–132 | EcoRI_*At*CNIH5.for | aagaattcATGGGCGATCTACTCGATTGGATTA |
| | *At*CNIH5 1–132_EcoRI.rev | aagaattcTAATGCCGAGTGGATCAACCAATA |
| NubG-*At*CNIH5 1–92 | EcoRI_*At*CNIH5.for | aagaattcATGGGCGATCTACTCGATTGGATTA |
| | *At*CNIH5 1–92_EcoRI .rev | gaattcCAAATGTTCCCTCTTCATGTATAGTCTAATGTTGT |
| NubG-*At*CNIH5 1–49 | EcoRI_*At*CNIH5.for | aagaattcATGGGCGATCTACTCGATTGGATTA |
| | *At*CNIH5 1–49_EcoRI .rev | gaattcCATTCGGTTTATTCTGCTAGATACATCGTAAGGG |
| NubG-*At*CNIH5 1–26 | EcoRI_*At*CNIH5.for | aagaattcATGGGCGATCTACTCGATTGGATTA |
| | *At*CNIH5 1–26_EcoRI .rev | aagaattcTGTCAGCTGATAAATAACGATGATTATAAG |
| **Plant binary vectors** | | |
| *At*SOS-S11 | BamHI_*At*SOS.for | ggatccATGGAGACGGAAACGAATCAGGT |
| | *At*SOS (noSTOP)_SalI.rev | gtcgacCTCAGCATCCACAACCGTAACCG |
| *At*DTX21-S11 | PacI_*At*DTX21.for | ttaattaaATGGCCGGAGGAGGAGGAGAGCTCA |
| | *At*DTX21 (noSTOP)_SalI.rev | gtcgacTTCCTCTGATGATACTTGGTTTACGTCTC |
| *At*DTX35-S11 | PacI_*At*DTX35.for | ttaattaaATGGATCCGACGGCGCCGTTGCTTA |
| | *At*DTX35 (noSTOP)_SalI.rev | gtcgacCGCAAGTATATCCTTGGATGTCGTCTCAC |
| *At*URGT6-S11 | AscI_*At*URGT6.for | ggcgcgccATGGCTCCAGTGAGTAAAGCTGAT |
| | *At*URGT6 (noSTOP)_SalI.rev | gtcgacGGCTTTGTCTTCGTTGTCGTCAG |
| *At*OCT1-S11 | SmaI_OCT1 CDS.for | cccgggATGGAACCTTCAAAACAAGAAGTTC |
| | OCT1_no stop_ XhoI.rev | ctcgagAGTAATCATGATTGTTTCGTTTTCG |
| *At*VPE1-S11 | AscI_VPE1 CDS.for | ggcgcgccATGGGTTCTCTAATGCAATCTGAAC |
| | VPE1 CDS_no stop_SalI.rev | GTCGACCACTTCCATCACATGATCTTGGACA |
| *At*HAK5-S11 | AscI_HAK5 CDS.for | ggcgcgccATGGATGGTGAGGAACATCAAATAG |
| | HAK5 CDS_no stop_SalI.rev | GTCGACTAACTCATAGGTCATGCCAACCTG |
| *AT*SUITR1;3-S11 | SmaI_SULTR1;3 CDS.for | cccgggATGTCGGCTAGAGCTCATCCTGTGG |
| | SULTR1;3_no stop_ XhoI.rev | ctcgagGACCTCGTCGGACAGTTTAGGGGAG |
| S10-*At*CNIH5 | AscI_*At*CNIH5.for | ggcgcgccaATGGGCGATCTACTCGATTGGATTA |
| | *At*CNIH5 STOP_XhoI.rev | ctcgagTTAGATGTCTCCTAATGCCGAGTGGA |

| | | |
|---|---|---|
| *At*PHT1;1-S11 | *At*PHT1;1 CDS XbaI.for | tctagaATGGCCGAACAACAACTAG |
| | *At*PHT1;1 CDS noSTOP SpeI.rev | actagtTTTCTCGTCATGGCTAACCTC |
